# AtCIPK16 Mediates Salt Stress Through Phytohormones and Transcription Factors

**DOI:** 10.1101/2020.02.17.953216

**Authors:** Shanika L. Amarasinghe, Wenmian Huang, Nathan S. Watson-Haigh, Matthew Gilliham, Stuart J. Roy, Ute Baumann

## Abstract

Soil salinity causes large productivity losses for agriculture worldwide. “Next-generation crops” that can tolerate salt stress are required for the sustainability of global food production. Previous research in *Arabidopsis thaliana* aimed at uncovering novel factors underpinning improved plant salinity tolerance identified the protein kinase AtCIPK16. Overexpression of *AtCIPK16* enhanced shoot Na^+^ exclusion and increased biomass in both Arabidopsis and barley. Here, a comparative transcriptomic study on Arabidopsis lines expressing *AtCIPK16* was conducted in the presence and absence of salt stress, using an RNA-Seq approach, complemented by AtCIPK16 interaction and localisation studies. We are now able to provide evidence for AtCIPK16 activity in the nucleus. Moreover, the results manifest the involvement of a transcription factor, AtTZF1, phytohormones and the ability to quickly reach homeostasis as components important for improving salinity tolerance in transgenics overexpressing *AtCIPK16*. Furthermore, we suggest the possibility of both biotic and abiotic tolerance through AtCIPK16, and propose a model for the salt tolerance pathway elicited through AtCIPK16.

## 1 Introduction

Soil salinity has adverse effects on global agricultural production (FAOSTAT 2014; Rengasamy 2010; Rengasamy 2006). An estimated 30% of the irrigated land and 6% of the world’s total land is affected by salt, and these areas are increasing in size (Schroeder et al. 2013). Estimates put agricultural production losses at 12 billion USD per annum in the US alone (Munns and Gilliham 2015; Shabala 2013). Finding crops that can withstand high salinity therefore is a high-priority for achieving sustainable world food production. Salinity imposes two main limitations on plant growth and survival: (i) an initial osmotic stress; and, (ii) secondary nutritional imbalance resulting in ionic and oxidative stress via accumulation of high concentrations of sodium (Na^+^) and chloride (Cl^−^) (Asif et al. 2019; Munns and Tester 2008; Roy et al. 2014; Shabala et al. 2015; Wungrampha et al. 2018). There is extensive research efforts focused on understanding the molecular mechanisms that confer increased salt tolerance with the ultimate goal of developing more salt tolerant crops (Deinlein et al. 2014; Hanin et al. 2016; Munns and Gilliham 2015; Roy et al. 2014).

Molecular mechanisms involved in salt tolerance in plants, can be broadly classified into the following categories: a) transporters that can reduce influx, or increase efflux or compartmentalization of Na^+^/Cl^−^ ions, or maintain K^+^ homeostasis (e.g. Salt Overly Sensitive 1 (SOS1), N^+^/H^+^ exchanger 1 (NHX1), High-affinity Potassium Transporter (HKT) 1;5, HKT1;4, HKT2;1, Potassium channel 1 (AKT1), Potassium transporter 5 (HAK5), Potassium channel KAT1, Cation-chloride cotransporter 1 (CCC1), S-type anion channel (SLAH1)) (Bassil et al. 2012; Chen et al. 2007; Diédhiou and Golldack 2006; Grabov 2007; Hamamoto et al. 2015; James et al. 2011; Ji et al. 2013; Mian et al. 2011; Qiu et al. 2016; Schachtman et al. 1992; Wang et al. 2015); b) detoxifiers that can scavenge excessive reactive oxygen species (ROS) and alleviate negative effects of ROS (e.g. Superoxide dismutases; SODs, ascorbate peroxidase; APX, ascorbic acid; AsA, catalases; CATs, GSH/glutathione peroxidase; GPX, NADH:peroxiredoxin oxidoreductase; PrxR) (Baxter et al. 2014; Mittler et al. 2011); c) osmotic adjusters that can maintain low intracellular osmotic potential in plants under salt stress (for example proline, glycine betaine, free amino acids, sugars, polyamines and polyphenols) (Rosa et al. 2009); d) phytohormones that can facilitate a broad array of adaptive responses and long distance signalling (such as abscisic acid, indole acetic acid, cytokinins, gibberellic acid, salicylic acid, brassinosteroids, jasmonates, ethylene) (Fahad et al. 2015; Peleg and Blumwald 2011; Ryu and Cho 2015); and e) salt sensors including those that sense cytosolic Ca^2+^ changes resulting from changes in the cytosol due to salinity and communicate the effects to downstream activating proteins (calcineurin-B-like proteins; CBLs and CBL-interacting protein kinases; CIPKs, calcium-dependent protein kinases; CDPKs, calmodulins; CaMs, calmodulin-like proteins; CAMLs, etc.) (Shabala et al. 2015).

The involvement of CBL-CIPK complexes as signalling components in salt stress has been well established (Batistič and Kudla 2009; Hashimoto et al. 2012; Luan 2009; Mao et al. 2016; Thoday-Kennedy et al. 2015). Arabidopsis CIPKs found to be involved in salinity tolerance mechanisms of plants include *CIPK1* (D’Angelo et al. 2006), *CIPK3* (Kim 2003), *CIPK6* (Tripathi et al. 2009), *CIPK16* (Roy et al. 2013) and *CIPK24* (*SOS2)* (Liu et al. 2000). *AtCIPK16* from *Arabidopsis thaliana* (*At2g25090*) was identified from a forward genetic screen as a gene with a role in reducing Na^+^ content in leaves during salt stress (Roy et al. 2013). Therefore, *AtCIPK16* is a potential candidate for the genetic engineering of salinity tolerant crops. The knowledge on the mode of action of AtCIPK16 is still largely unknown; however, previous studies have shown that AtCIPK16 has a nuclear localisation signal (NLS) (Amarasinghe et al. 2016).

The current study is an attempt to fill a gap in our understanding of *AtCIPK16* mediated salt stress tolerance in *A. thaliana.* Through an investigation of the transcriptomic responses in transgenic and null-transgenic plants, as well as a co-expression network analysis, we aimed to identify a set of genes, whose expression is influenced directly or indirectly by *AtCIPK16* overexpression. Our results suggest that the *AtCIPK16* mediated salt tolerance is mainly achieved through transcription factor modulation and phytohormone signalling. We propose a molecular pathway for at least a part of the *AtCIPK16* mediated salt tolerance mechanism for validation in future laboratory experiments.

## 2 Materials and Methods

### 2.1 Yeast two hybrid-assay

Full-length of the coding sequences of all 10 AtCBLs and AtCIPK16 were amplified from cDNA of Arabidopsis ecotype Columbia-0 by high fidelity PCR using primers listed in Supp Table 1 and cloned into a pCR8/GW/TOPO TA Gateway® entry vector (Invitrogen, USA) following the manufactures protocols. An LR reaction was performed to transfer the DNA from pCR8 into the required destination vectors, the prey vector pTOOL28 and the bait vector pTOOL27. The AtCIPK16 bait vector, 10 AtCBLs prey vectors were co-transformed into the yeast strain AH109 following manufacturer’s instructions (Clontech, CA, USA). The empty vectors pTOOL27 and pTOOL28 were also co-transformed into yeast with each prey or bait plasmid respectively to confirm that bait does not autonomously activate the reporter genes in the absence of the prey protein.

**Table 1.**
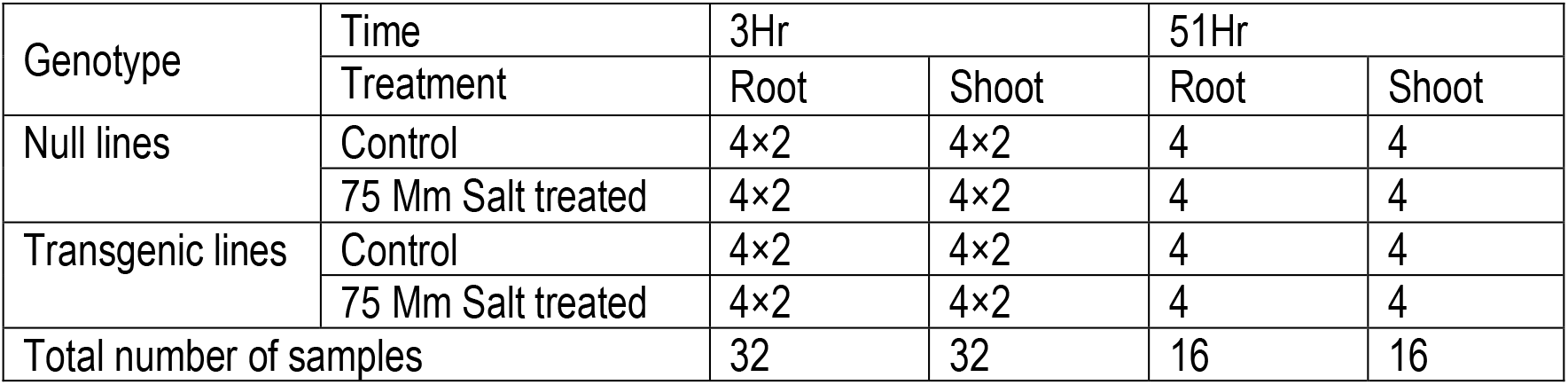
Experimental design for the current study. There were 4 biological replicates per condition per experimental group for both time points. The 3 hour samples were sequenced in two technical replicates per biological sample hence indicated as a multiple of 2.

| Genotype | Time | 3Hr |  | 51Hr |  |
| --- | --- | --- | --- | --- | --- |
|  | Treatment | Root | Shoot | Root | Shoot |
| Null lines | Control | 4×2 | 4×2 | 4 | 4 |
|  | 75 Mm Salt treated | 4×2 | 4×2 | 4 | 4 |
| Transgenic lines | Control | 4×2 | 4×2 | 4 | 4 |
|  | 75 Mm Salt treated | 4×2 | 4×2 | 4 | 4 |
| Total number of samples |  | 32 | 32 | 16 | 16 |

For each possible combination between bait and preys the largest colonies were picked from each plate and inoculated into 5 mL selection SD QDO liquid medium (containing 2 % glucose), followed by incubation at 30 °C with shaking at 180 rpm (orbital mixer incubator, Ratek Instruments, Australia) for 2 days. OD_600_ of yeast liquid samples were determined by an ND-1000 spectrophotometer (NanoDrop Technologies Inc.) and adjusted on OD_600_ of 0.25 with sterile Milli Q water. To establish whether the bait and prey proteins interacted on the plates, 10 μL of 10-fold serial dilutions of yeast harbouring an activation domain (AD)-AtCBLs or empty vector pTOOL28 and either DNA-binding domain (BD)-AtCIPK16 or empty vector pTOOL27 were spotted on plates containing DDO (SD/-Leu/-Trp, as a control) and QDO (SD/-Leu/-Trp/-His/-Ade (Clonetech) to determine whether there is an interaction between prey and bait molecules. Both DDO and QDO selection plates were incubated at 30 °C for 5-8 days to prior to the growth of yeast being observed.

### 2.2 Bimolecular fluorescence complementation (BiFC) assay using both transient expression and stable expression

To confirm the interaction of AtCIPK16 with 10 AtCBL proteins and to localise the interaction complex of AtCBL-AtCIPK16 in Planta, a bimolecular fluorescent complementation (BiFC) assay was performed using transient expression in tobacco leaf epidermal cells.

The full-length coding sequence of 10 AtCBLs and AtCIPK16 in pCR8 with or without the stop codon were cloned into a pCR8/GW/TOPO TA Gateway® entry vector (Invitrogen, USA). An LR reaction was performed to transfer genes into a pGPTVII (need ref) destination vector. Consequently, each AtCBL was expressed as a fusion to C-terminal YFP fragment (YC) and AtCIPK16 was expressed as a fusion to N-terminal YFP fragment (YN). Constitutive expression of the genes of interest in plants is ensured by the 35S promoter of the cauliflowers mosaic virus in the vectors.

Pairs of 10 AtCBLs and AtCIPK16 in pGPTVII constructs were co-transformed into A.tumefaciens AGL1 cells using the protocol adapted from Kapila et al.(1997) and Wydro et al. (2006). Mixtures containing pairs of AtCBLs-AtCIPK16 were aspirated into a 1 mL needless syringe (Terumo, Tokyo, Japan) and injected into the abaxial side of the third and fourth leaves of different six-week-old tobacco until saturated. After infiltration, the transformed tobacco plants were then kept in short day growth room (light/dark period: 10h/14h, temperature: 23 °C, light intensity: 100 μmol m-2 s-1, 60-75 % humidity) for 3-4 days to allow the expression of AtCBLs-AtCIPK16 before harvesting leaf material for visualisation by confocal.

Location (if any) of the interaction of fluorescently tagged AtCIPK16 and the different AtCBLs in leaf cells was determined using a A Zeiss Axioskop 2 MOT plus LSM5 PASCAL confocal laser scanning microscopy equipped with an argon laser (Carl Zeiss, Jena, Germany).: YFP signal was detected using an excitation of 514 nm and an emission bandpass of 570-590 nm. Propidium iodide/chlorophyll autofluorescence was detected using an excitation of 543 nm and an emission longpass filter of 560 nm. False colour images were analysed using LSM5 Image Examiner (Carl Zeiss, Jena, Germany).

### 2.3 RNA-Seq Assay

#### 2.3.1 Experimental Design

The RNA-Seq study has a 2 (genotype: null, transgenic) by 2 (tissue: root, shoot) by 2 (treatment: control, salt treated) by 2 (time: 3 hr, 51 hr) factorial design. To ensure a minimum number of 4 biological replicates for the RNASeq analysis six *A. thaliana* replicate plants were sampled for each experimental group of which 4 were sent for sequencing. The final experimental design is summarised in Table 1. For the 3 hour time point there were two technical replicates per biological sample hence represented as a multiple of 2.

#### 2.3.2 Transformation of *AtCIPK16*, T2 seed germination, Plant material, growth conditions, salt treatment and sampling

Transgenic *35S:AtCIPK16* overexpressing *Arabidopsis thaliana*, Col-0, were previously generated as described in Roy et al. 2013. The plant growth in hydroponics was conducted according to Jha et al. (2010). Seeds of T_2_ *35S:AtCIPK16* plants were soaked in 70% ethanol for 2 minutes followed by 4-5 rinses in sterile milli-Q water for surface sterilisation. Subsequently the seeds were planted in 1.5 mL microfuge tubes containing half-strength *Arabidopsis* nutrient solution (Arteca and Arteca 2000) and 0.8% Bacto agar. After vernalisation for 2 days at 4°C, seeds were transferred to a growth room with controlled light (10 hr light/14 hr photoperiod, an irradiance of 70 mmol m^−2^s^−1^) and constant temperature of 21 °C. After emergence of the seedling roots, plants were transferred to a hydroponic tank containing full-strength *Arabidopsis* nutrient solution (Arteca and Arteca 2000). The pH of the hydroponic solution was maintained at 5.7. After 5 weeks of growth in hydroponics, salt stress was applied to half the plants by the addition of 75 mM NaCl. Calcium activity in the growth medium was maintained at 0.3 mM by the addition of the correct amount of CaCl_2_, as calculated using Visual Minteq Version 2.3 (KTH, Department of Land and Water Resources Engineering, Stockholm, Sweden). Shoot and root tissues were removed after 3 hours and 51 hours of salt stress for RNA extraction and were immediately frozen in liquid nitrogen. To avoid effects on gene expression due to the circadian rhythm the second sampling was conducted at 51 hours rather than 48 hours (2 days) after stress treatment RNA isolation, library preparation and Illumina sequencing. Prescence of the transgene in the *AtCIPK16* expressing lines was determined as before (Roy et al., 2013).

Total RNA was extracted using the TRIzol reagent (Invitrogen, Carlsbad, CA, USA), following the protocol described by Chomczynski (1993). TruSeq™ stranded RNA sample preparation was utilized with dUTP second strand marking protocol for cDNA library preparation. Ribo-Zero kit (Epicentre, an Illumina company, Madison, WI) was used to remove rRNA from the libraries. Illumina Sequencing was carried out to collect 100bp paired-end (2 * 100bp). Aim was to get a read depth of 15 Mill read pairs per library.

#### 2.3.3 RNA-Seq data pre-processing

Raw data was examined by the program FASTQC for read quality, detection of adapter contaminations and presence of overrepresented sequences (Andrews 2010). Next, a Java based in-house *k-mer* counting algorithm was used to confirm the presence/absence of the transgene in each sample by counting reads belonging to the UTRs of the transgene and the wild type respectively. As described in Roy et al. (2013), the *CIPK16* transgene was inserted inbetween the 35S CaMV promoter and 3^′^UTR of the *nos* terminator. To distinguish expression of the transgene from the endogenous *AtCIPK16* in Col-0 plants, an k-*mer* counting script available in LNISKS package (https://github.com/rsuchecki/LNISKS; Suchecki et al., 2019) was supplied with these sequences (*AtCIPK16* 5^′^ UTR, *AtCIPK16* 3^′^ UTR and region in between the *AtCIPK16* exon 1 and the *AtCIPK16* 5′ UTR) to count reads belonging to the expressed transgene from the FASTQ files. Furthermore, regions 20 kb upstream and downstream of the *AtCIPK16* gene obtained from the TAIR database (https://www.arabidopsis.org/) were used to count the reads expressed from the endogenous wild-type *AtCIPK16*.

Reads with lengths ≥ 70 bp after quality trimming were used for further processing. The Arabidopsis reference genome TAIR10, and gene model annotation files were downloaded from the TAIR ftp site (http://www.tair.org). Read alignment to the reference genome was performed using the splice aligner STAR (version 2.4.1c) (Dobin et al. 2013). There are two steps in mapping using the STAR aligner. 1; Building a genome index for the reference genome (FASTA sequences): *A. thaliana* GFF file was used with an overhang of 100 (i.e. max *readLength* −1) for creating the index. The chloroplast and mitochondrial genomes were excluded when creating these index files. 2; Mapping the reads to the genome: the paired-end reads were mapped to the reference with no mismatches allowed (because both reference and samples were from Col-0 cultivar), with a maximum intron size of 2000 and a maximum gap of 2000 between two mates. Alignment results were output in Sequence/Binary Alignment Map (SAM/BAM) format sorted by the chromosome coordinates. Alignments with non-canonical junctions were filtered out. Quantification of gene expression level and identification of Differentially Expressed Genes (DEGs)

Read counting for the transcripts was done using the *featureCounts()* function of the *Rsubread* package (Liao et al. 2014) implemented in the R environment (http://www.R-project.org). *DGEList()* function from *DESeq* R library was used to calculate the counts per million (CPM) for each experimental group based on the count matrix, and the *calcNormFactors()* function was applied to estimate normalization factors (Anders and Huber 2010). Data from each time point (i.e. 3 hr, 51 hr) and each tissue were analysed separately resulting in 4 groups (i.e. Root_3Hr, Shoot_3Hr, Root_51Hr and Shoot_51Hr). Transcripts with CPM values of less than 100 in 75% of the samples or more were filtered out. Out of 64, 60 samples at the 3 hour time point were analysed. A sample from root and the corresponding shoot sample (of which each had 2 technical replicates; Table 1) from the 3 hour time point had to be removed due to a large variation in *AtCIPK16* expression compared to other samples (S3).

T-statistics for mean expression values for each gene was determined using the LIMMA (Linear Models for Microarray Data) package implemented in the R software environment (Ritchie et al. 2015; Smyth 2004). The read count data fed into the Limma linear model fitting were transformed using *Voom* with quantile normalization followed by group-means parameterization and robust eBayes (Law et al. 2014; Smyth 2004). The contrast matrix was created with the final goal of; identifying the differentially expressed genes in salt stress that is due to the definite effect of the transgene. This contrast matrix was used on the 4 groups of expression value data separately. P-values for multiple testing were corrected according to Benjamini and Hochberg 1995. Differentially expressed genes are those with a FDR-adjusted P value of ≤0.05 and ≥2-fold change in expression relative to the control.

#### 2.3.4 Regulatory network construction using WGCNA

Weighted Gene Co-expression Network Analysis (WGCNA) enables the detection of modules of genes with high expression value correlation to one another (Langfelder and Horvath 2008). Briefly, WGCNA assigns a connection weight between pairs of genes within the network based on a biologically motivated criterion and attempts to identify relevant modules by applying a soft threshold to correlations between pairs of genes within a network. R software environment was used for all WGCNA analysis (Langfelder and Horvath 2008; Zhang and Horvath 2005). After confirming that there were no outliers in the tissue-separated samples optimization of the soft threshold values was performed. Signed co-expression network was constructed using the automatic one-step network construction method (function *cuttreeDynamic()*) with following settings; a signed type of network, an unsigned type of topological overlap matrix (TOM), correlations of the network raised to a soft thresholding power β (roots:10; shoots: 5), correlation measures with option ‘*bicor*’, deepSplit value of 2, a minimum module size of 20. The first principle component of a module (module eigengene) value was calculated and used to test the association of modules with salt response in the null and transgenic genotypes. Total network connectivity (kTotal), and module membership (MM), were calculated for each of the DEGs. Modules for further analysis were selected if one or more of their hub genes (genes with modular membership ≥ 0.9) was among the transgene dependent salt-responsive gene list of the respective tissue.

#### 2.3.5 Functional analysis

GO term enrichment was performed using the Plant Gene Set Enrichment Analysis Toolkit (PlantGSEA) and the DAVID online web server (http://david.abcc.ncifcrf.gov/) (Dennis et al. 2003; Yi et al. 2013). Up and down regulated DEGs from each contrast were used for GO and pathway enrichment analysis, and a False Discovery Ratio (FDR) corrected p value ≤ 0.05 was selected as the threshold level of significance to determine the enrichment in the gene set. MapMan stand-alone software allowed the assignment of DEGs into regulatory pathways (Thimm et al. 2004). Additionally, Kyoto Encyclopaedia of Genes and Genomes (KEGG) was used to identify higher order functional information related to the DEGs (Kanehisa et al. 2017). ATTED-II (http://atted.jp) gene co-expression database was mined to identify additional yet relevant genes that may be co-expressed with the DEGs (Aoki et al. 2016).

To identify genes that are exclusively differentially expressed only in transgene dependent manner in presence of salt, the DEGs from the transgene dependent salt responsive gene list was compared in a pairwise manner to the DEGs from that of transgene effect in controls for a given tissue at a particular time point.

Phosphorylation targets were identified using the NetPhoS4.1 online server (Blom et al. 1999). MPK substrates were identified by comparing the DEGs to known substrates recorded in the literature (Meng et al. 2013; Nguyen et al. 2012; Popescu et al. 2009; Vogel et al. 2012). Promoter analysis for the ARE motif; regions 3000 bp upstream from the transcription start site (TSS) of all transgene dependent salt responsive genes of root and shoot at 3 hours were downloaded from the TAIR database (https://www.arabidopsis.org/). The motif pattern ATTTATTTATTT{A|T] was searched against the downloaded sequences through the FIMO tool (http://meme-suite.org/tools/fimo) in the MEME suite (Grant et al. 2011). The p-value threshold was set to 0.01. DNA binding domains and amino acid properties were identified from protein sequences using the consensus of results obtained through several freely available online tools with the use of their default settings; DP-BIND (Hwang et al. 2007), BINDN (Wang and Brown 2006), NetSurfP (Petersen et al. 2009), PredictProtein (Rost et al. 2004), paircoil2 (McDonnell et al. 2006) and pepinfo (Li et al. 2015).

### 3 Results

### 3.1 Determining AtCIPK16 interacting CBLs

AtCIPK16 was found to interact with 6 of the 10 AtCBL proteins. AtCIPK16 was found to interact strongly with AtCBL4 and AtCBL5; moderately with AtCBL2 and AtCBL9; and weakly with AtCBL1 and AtCBL10 (Figure 1). In contrast, AtCIPK16 did not interact with AtCBL3, AtCBL6, AtCBL7 and AtCBL8 (Figure 1). Similar growth of all yeast samples were observed on the control SD (-Leu/-Trp) plates (Figure 1).

**Figure 1.**
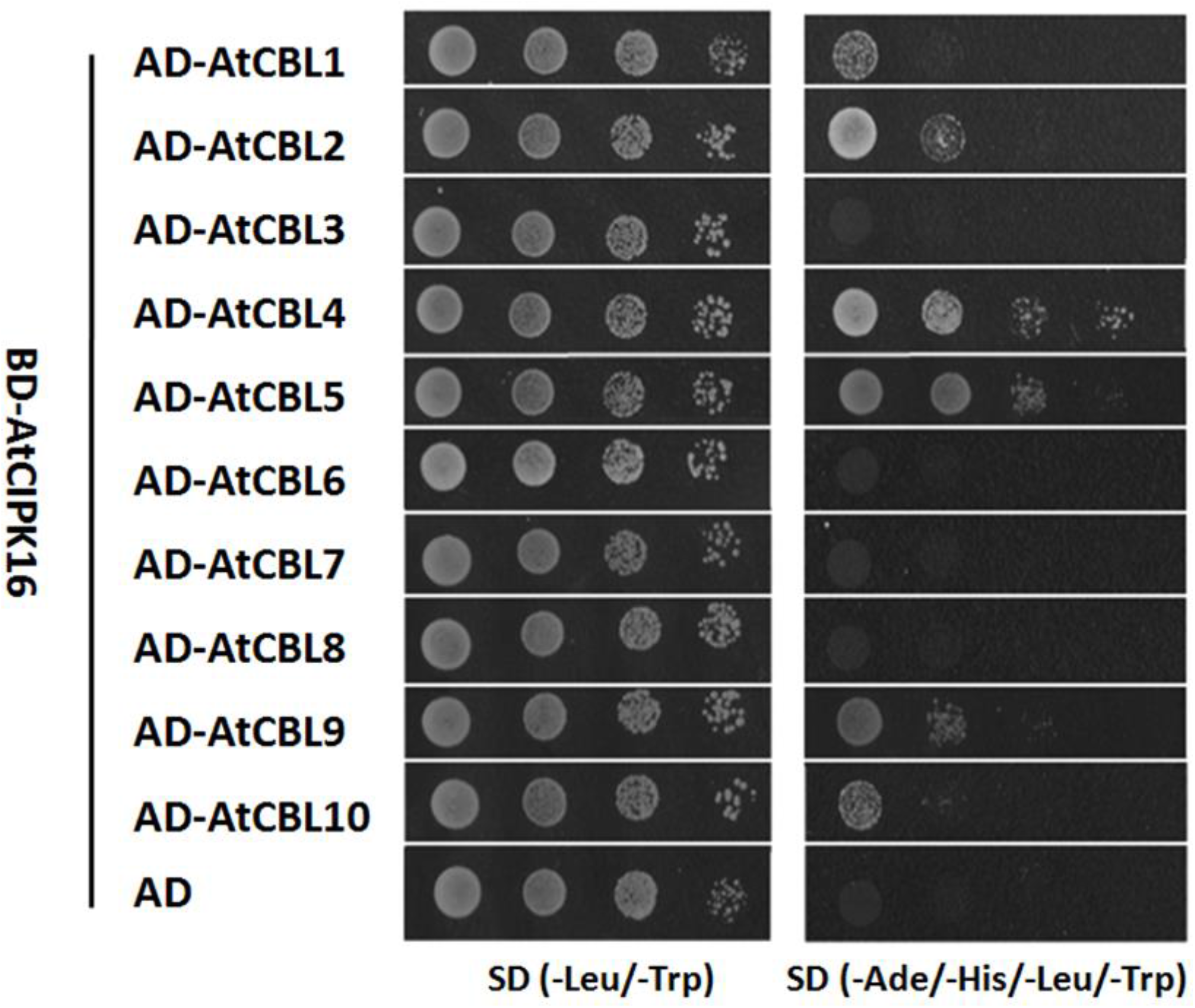
AtCIPK16 interacts with 6 AtCBL proteins. 10-fold serial dilutions (OD_600_=1, 10-1, 10-2, 10-3) of yeast sample harbouring a DNA-binding domain (BD)-AtCIPK16 and either an activation domain (AD)-AtCBLs or an empty vector (AD) were grown on selection DDO (SD/-Leu/-Trp) and QDO (SD/-Leu/-Trp/-His/-Ade) media. Both DDO and QDO selection plates were incubated at 30 °C for 6 days to allow the yeast to grow.

### 3.2 Resolving subcellular localization of AtCIPK16:CBL interactions

YFP fluorescence in the nucleus and possibly at the plasma membrane was observed for AtCBL1-AtCIPK16, AtCBL4-AtCIPK16, AtCBL5-AtCIPK16 and AtCBL9-AtCIPK16 (Figure 2 a, d, e and i). AtCBL2-AtCIPK16 and AtCBL10-AtCIPK16 complexes were observed in the cytoplasm and nucleus (Figure 2 b and j), while AtCBL3-AtCIPK16 was observed in the cytoplasm (Figure 2c). However, no clear fluorescent signal was observed for AtCBL6-AtCIPK16, AtCBL7-AtCIPK16 and AtCBL8-AtCIPK16 complexes (Figure 2 f, g and h). Based on this observation, AtCIPK16 sequence was tested for DNA binding potential. It was predicted that the protein region from A^357^ – G^391^ has the ability to bind to DNA (Figure 3).

**Figure 2.**
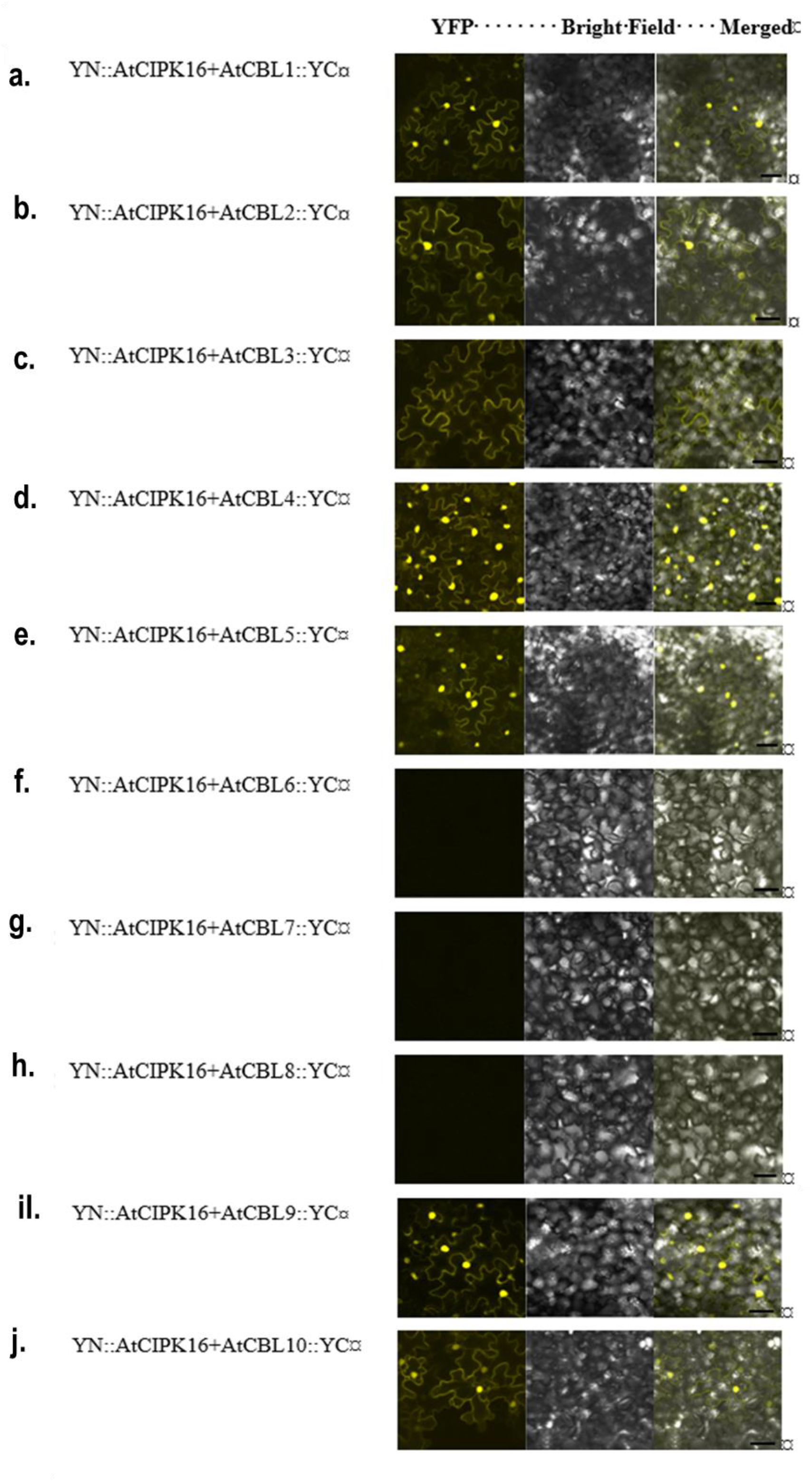
YN::AtCIPK16 and AtCBLs::YC interactions in tobacco leaves (*Nicotiana benthamiana*) Images of tobacco leaf with co-infiltration of different pairs of YN::AtCIPK16 and AtCBLs::YC constructs. Leaves were visualised using confocal microscopy and YFP fluorescence was captured using the wavelengths: excitation, 514 nm; emission, 525–538 nm. (A-J) Images of tobacco leaves with co-transformed with each pair of AtCBLs::YC and YN::AtCIPK16. The images in the first column show YFP fluorescence in leaves, the images in the second column show the bright field of leaves and the third column is a merged image of the YFP fluorescence and bright field of leaves. Scale bar = 50 µM.

**Figure 3.**
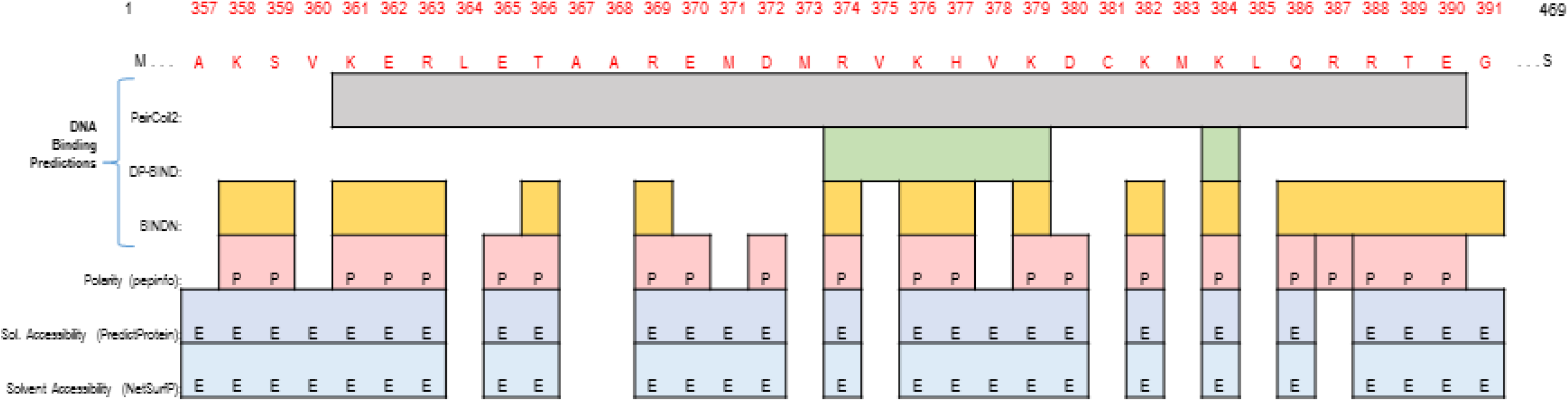
Identified region of AtCIPK16 protein with DNA binding affinity. The amino acid residue numbers and the respective residues are mentioned in the first and second rows, respectively; residues in the putative region with DNA binding ability are shown in red. The server result summary for each residue is shown below the respective amino acid. Row 3-5 show the predicted DNA binding affinity by three independent online tools (i.e. PairCoil2; grey, DP-BIND; green and BINDN; yellow); 6th row shows the polarity prediction for the region using pepinfo server (denoted with a P in red background); 7th and 8th rows show the solvent accessibility predicted by two independent servers (PredictProtein and NetSurfP, respectively; denoted by E in blue background.

### 3.3 Determining presence and transgene expression level in *35S:AtCIPK16* expressing Arabidopsis

Presence of the transgene was determined by using primers designed to the transgene specific 3^′^ UTR region of the gene and confirmed only transgenic plants contained the *AtCIPK16* transgene (S1). *AtCIPK16* transgene expression was higher in transgenic plants compared to native *AtCIPK16* expression in null segregants in the absence of salt stress, and after both 3 and 51 hours of salt stress (Figure 4).

**Figure 4.**
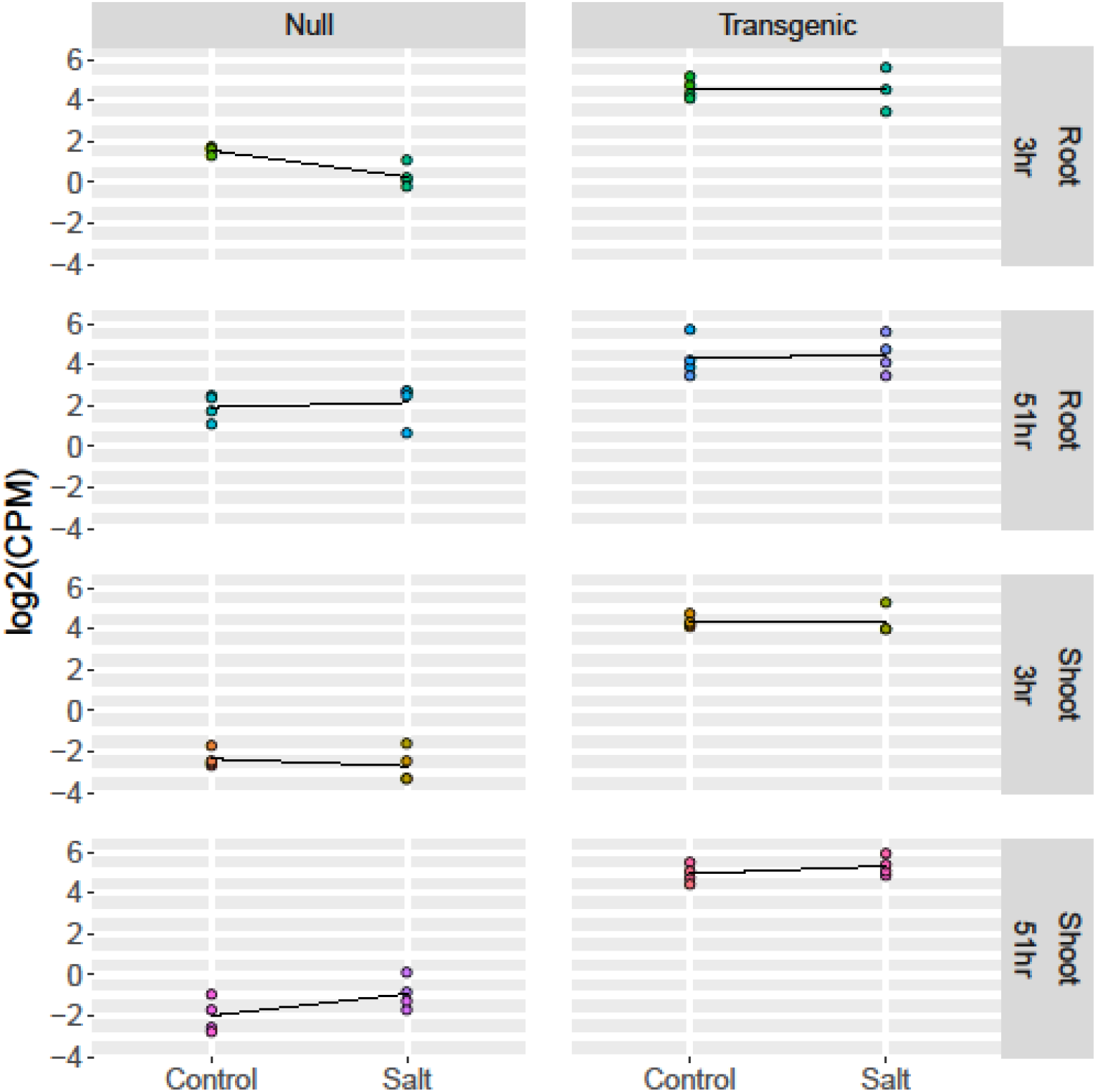
*AtCIPK16* gene expression in the current study. The expression values are measured in counts per million (CPM) and displayed as log_2_(CPM) for clarity (y axis). Each dot represent a sample and is coloured according to the experimental condition. Expression values are separated by genotype (i.e. null, transgenic: on top) and tissue-time point (i.e. root 3hr, shoot 3hr, root 51 hr and shoot 51hr: right) and treatment (i.e. control, salt: bottom). A black solid line connects the mean expression value from the two treatments in each group of samples. A gene is considered expressed if the mean of log expression value is above 0 for a given experimental condition.

### 3.4 Differential Gene Expression

To determine differential gene expression between *AtCIPK16* over-expressing lines and null segregants, RNA was extracted from shoot and root material of 5 week old, hydroponically grown, plants exposed to either 0 or 75 mM NaCl for 3 or 51 hours. RNA-Seq analysis was performed to determine the plants’ gene expression profiles. On average, a mapping percentage of ∼88% was reported across root and shoot material collected from both transgenic and null segregants for the 3 hour time point data and a mapping percentage of ∼86% for 51 hour time point samples (S2).

A total of 21,974 and 21,160 genes were differentially expressed across the two tissues from 3 hours and 51 hours, respectively, in salt treated *AtCIPK16* transgenic plants compared to the null transgenics. In order to identify the differentially expressed genes in salt stressed plants with *AtCIPK16* overexpression several contrasts were tested (Figure 5) based on the differences in gene expression levels in the roots at 3 hours, shoots at 3 hours, roots at 51 hours and shoots at 51 hours. The number of genes which were up or down-regulated at each of the two time points and in each tissue for each line are shown in Figure 6.

**Figure 5.**
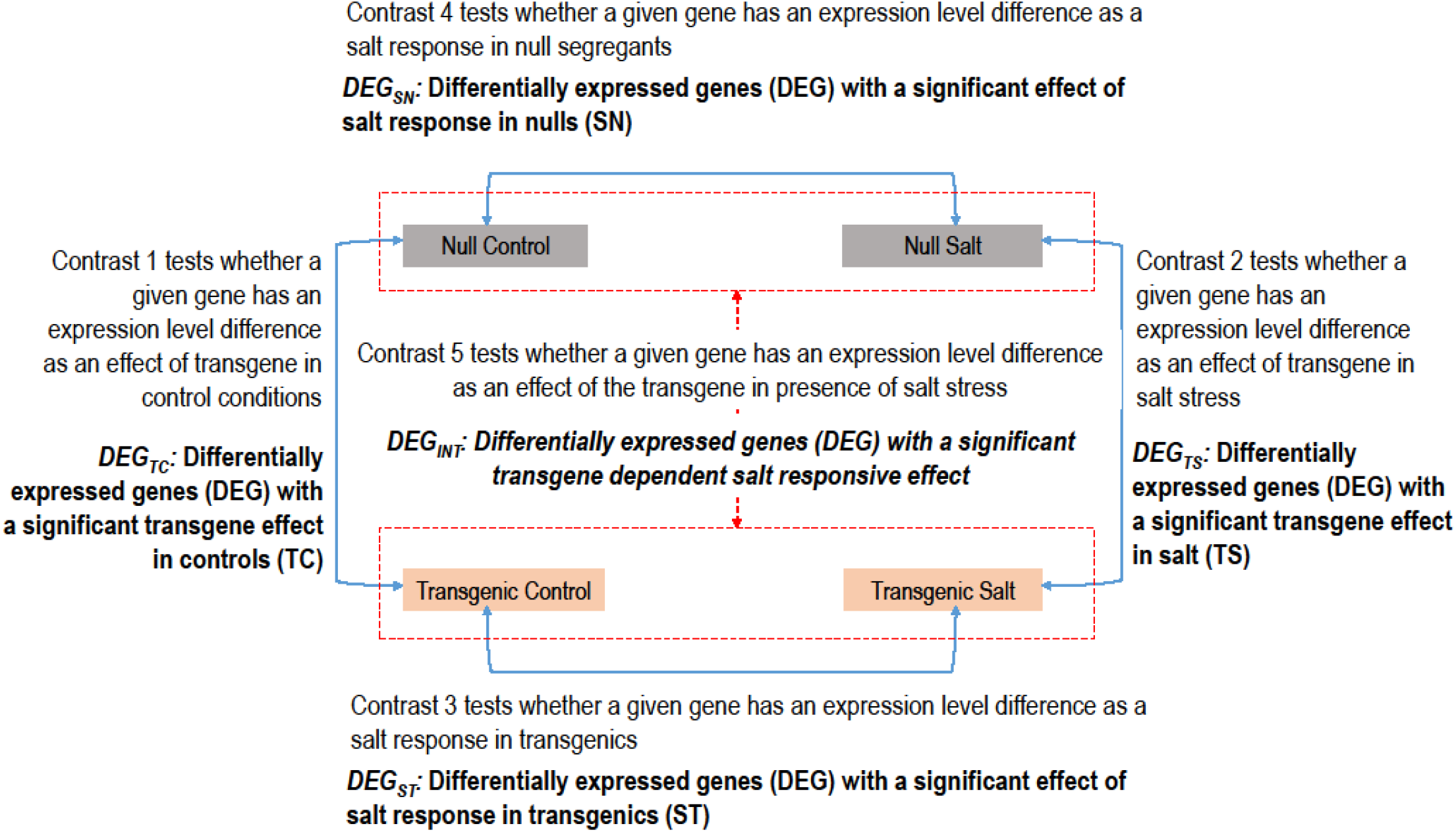
Contrasts tested in the current analysis. The experimental groups compared by Limma (Ritchie et al, 2015) to test each contrast are shown by blue two headed arrows. The red dashed boxes and two headed arrow denote the contrast to test the interaction term. The term that defines the differentially expressed genes for each contrast is mentioned below each hypothesis in ***bold italics***

**Figure 6.**
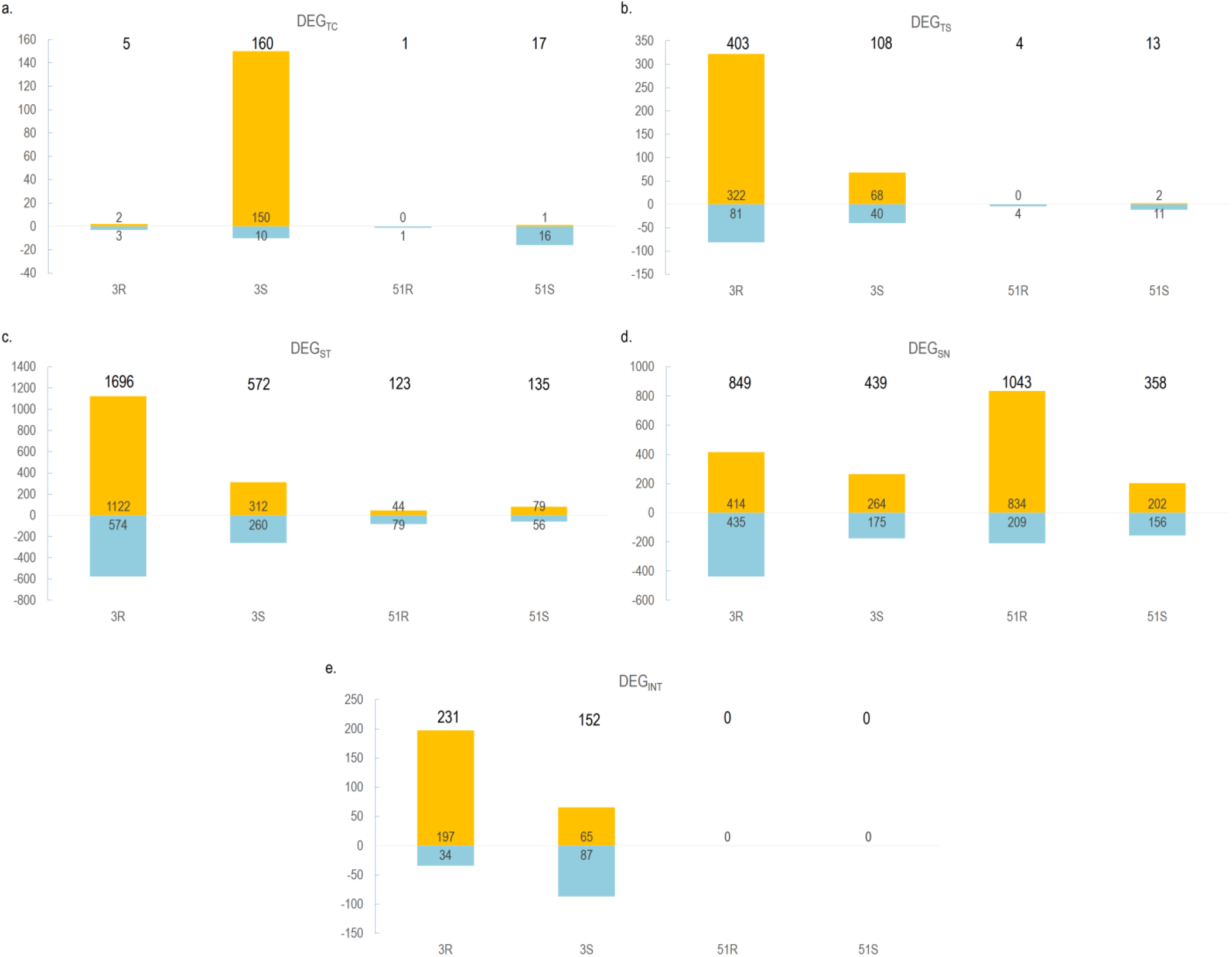
Number of genes differentially identified. The number of differentially expressed genes (DEG) for each test shown in Figure 5 are shown for; a) transgene effect in controls (DEG_TC_); b) transgene effect in salt (DEG_TS_); c) salt effect in transgenics (DEG_ST_); d) salt effect in nulls (DEG_SN_) and e) transgene dependent salt responsiveness (DEG_INT_). Y axis displays the number of DEG; x axis represent the experimental group (3R: 3 hr root; 3S: 3 hr shoot; 51R: 51 hr root; 51S: 51 hr shoot) through Limma analysis (Ritchie et al, 2015). Yellow denoted the upregulated genes and blue denoted the down regulated genes. The total number of DEGs are shown on top of each bar. Individual numbers for up and down regulated genes are shown along the y axis on yellow and blue bars

#### 3.4.1 Contrast 1: Transgenic Control Vs Null Control

In the comparison of transgenic control vs null control (transgene-effect in controls; TC) samples, 5 differentially expressed genes (DEG_TC_) in the roots at 3 hrs (DEG_TC(3R)_) and 160 in the shoots at 3 hrs (DEG_TC(3S)_) were identified (Figure 6a, S4 worksheet 1-2). As expected, *AtCIPK16* (*AT2G25090*) was present in both DEG_TC(3R)_ and DEG_TC(3S)_ (Figure 7a). Most (150) of the genes in DEG_TC(3S)_ had higher expression in transgenics.

At 51 hours, there was only 1 DEG_TC_ in roots (i.e. DEG_TC(51R)_) (Figure 6a; S4 worksheet 3). In shoot controls at 51 hours there were 17 DEG_TC_ (i.e. DEG_TC(51S)_) (Figure 6a; S4 worksheet 4). While there is 1 DEGs in common between the root and the shoot DEG_TC_ at 51 hours, it is not *AtCIPK16* but *AT1G47970* (Figure 7b).

**Figure 7.**
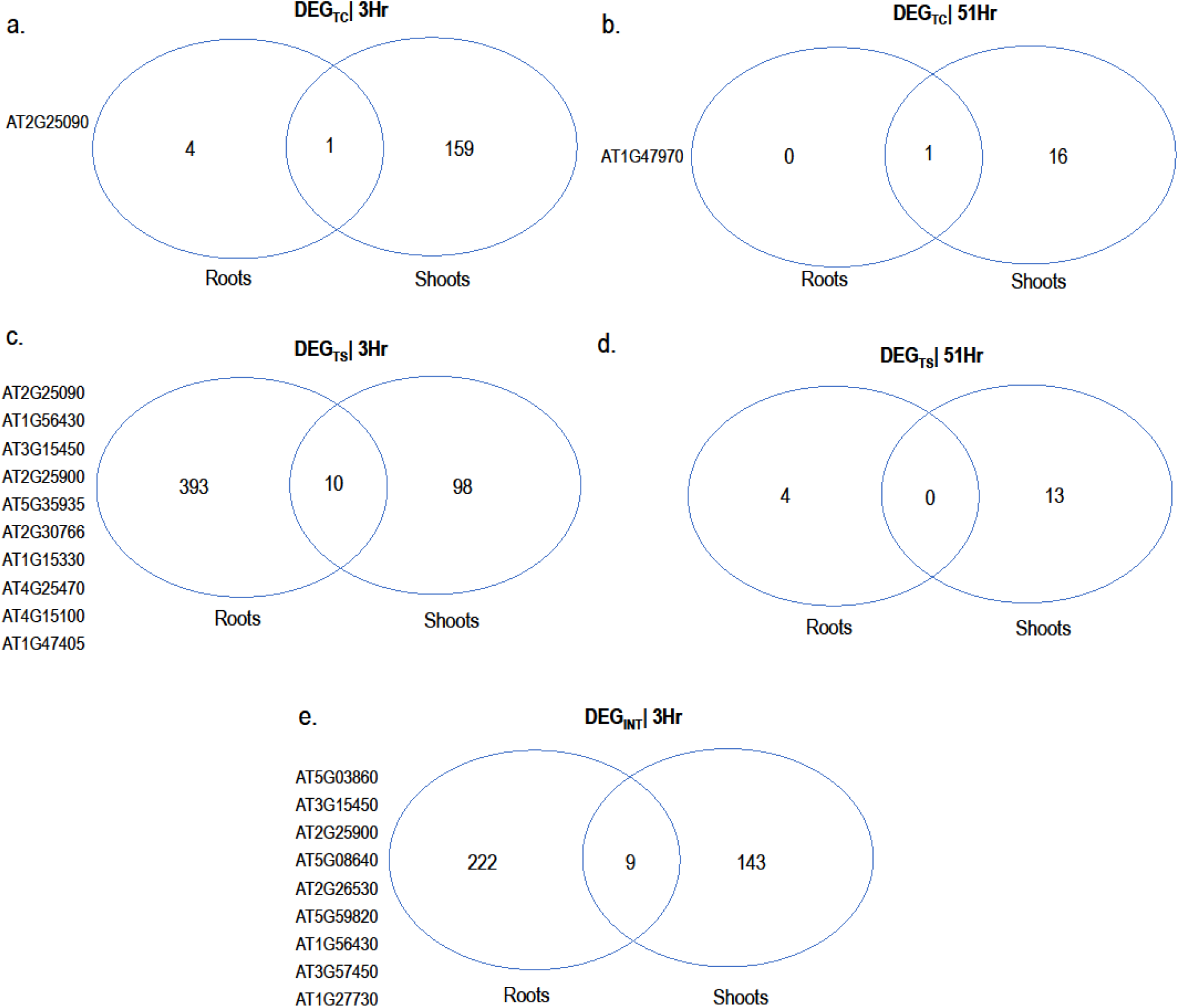
Comparison of the DEGs. Overlap of differentially expressed genes (DEG) in roots and shoots. The comparisons that were tested (Figure 5) are given at the top and the genes which overlap between the two tissues for that treatment are listed on the left side of each venn diagram. DEG_TC_: DEG from transgenic controls; DEG_TS_: DEG from transgenics in salt; DEG_TC_: DEG from transgene dependent salt responsiveness

#### 3.4.2 Contrast 2: Transgenic Salt Vs Null Salt

This is the comparison of transgenic vs null samples in presence of salt (transgene effect in salt: TS) (Figure 5) There were DEGs (DEG_TS_) present for this contrast for both tissues at both time points (i.e. DEG_TS(3R)_: 403, DEG_TS(3S)_: 108, DEG_TS(51R)_: 4, DEG_TS(51S)_: 13) (Figure 6b; S4 worksheet 5, 6, 7 and 8). While there was a ∼80 fold increase in the DEG_TS(3R)_ compared to DEG_TC(3R)_ (403 vs 5), the DEG_TS(3S)_ remained more or less in the same range as DEG_Tc(3S)_ (160 vs 108) (Figure 6 a and b). However, there were proportionally more down regulated genes in the DEG_TS(3S)_. While only ∼6% of the DEGs from DEG_TC(3S)_ were down regulated (10/160), ∼37% of DEG_TS(3S)_ were down regulated (40/108). Clearly during salt stress, overexpression of *AtCIPK16* reduced the expression of genes.

At 51 hours, a very low number of DEGs were seen in salt stress in both roots and shoots similar to the observations in non-stressed conditions (4 DEG_TS(51R)_ vs 1 DEG_TC(51R)_ and 13 DEG_TS(51S)_ vs 17 DEG_TC(51S)_) (Figure 6 a and b). There were 10 genes in common between DEG_TS_ of root and shoots at 3 hours (Figure 7c) while there were none at 51 hours (Figure 7d).

#### 3.4.3 Contrast 3: Transgenic Salt Vs Transgenic Control and Contrast 4: Null Salt Vs Null Control

At 3 hours, in both tissues the effect of salt (salt effect on transgenics: ST; Figure 5) elicited differential expression of more genes (DEG_ST_) in *AtCIPK16* transgenics (i.e. DEG_ST(3R):_ 1696 and DEG_ST(3S):_ 572) compared to the null transgenics (salt effect on nulls: SN; Figure 5) (DEG_SN(3R):_ 849 and DEG_SN(3S):_ 439) (Figure 6 c and d). But at 51 hours it is the opposite; effect of salt elicited the differential expression of fewer genes in *AtCIPK16* transgenics (i.e. DEG_ST(51R):_ 123 and DEG_ST(51S):_ 135) compared to nulls (DEG_SN(51R):_ 1043 and DEG_SN(51R):_ 358) (Figure 6 c and d).

#### 3.4.4 Contrast 5: Interaction between SN and ST

With the presence of DEGs in both ST and SN, genes with significantly different expression levels between these two contrasts were examined through linear modelling of data. The results of contrast 5 are differentially expressed genes due to the absolute effect of transgene in salt stress (INT) (i.e. DEG_INT_) (Figure 5). Even though for 3 hours there were 231 DEG_INT_ in roots (DEG_INT(3R)_) and 152 DEG_INT_ in shoots (DEG_INT(3S)_), there were no DEG_INT_ for the 51 hours (Figure 6e; S4 worksheet 9, 10, 11 and 12). Furthermore, there were 9 DEG_INT_ common to both roots and shoots at 3 hour time point (Figure 7e).

### 3.5 Investigating potential biological implications of *AtCIPK16* overexpression

Genes with a significant transgene-effect in controls (5 DEG_TC(3R)_, 160 DEG_TC(3S)_, 1 DEG_TC(51R)_ and 17 DEG_TC(51S)_) (Figure 6a) were further analysed through Gene Ontology (GO) studies and pathway analysis to understand potential biological consequences of *AtCIPK16* overexpression in the non-stressed conditions. GO analysis showed that DEG_TC(3S)_ that are up-regulated were most enriched for response to chitin (p value = 3.47×10^−92^) (S5; worksheet1 cells with yellow background colour). Perhaps not surprisingly, the corresponding Kyoto Encyclopaedia of Genes and Genomes (KEGG) pathways that DEG_TC(3S)_ fell into include plant-pathogen interaction (S6, column B). Additionally, molecular functions such as transcription regulator activity was significant for the DEG_TC(3S)_ that are up-regulated (p value = 7.51×10^−09^) (S5; worksheet 1 cells with yellow background colour). Down-regulated DEG_TC(3S)_ were enriched for the molecular function of negative regulation of RAS protein signal transduction (p value = 6.95×10^−04^) and RHO-GTPase binding (p value = 6.95×10^−04^) (S5, worksheet1 cells with blue background colour).

The significant GO terms found for up-regulated DEG_TS(3R)_ were the biological process response to organic substance (p value = 1.59×10^−42^) and the molecular function sequence specific DNA binding transcription factor activity (p value = 5.65×10^−11^) (S5, worksheet 2 cells with yellow background colour). The up-regulated DEG_TS(3S)_ were enriched for cell wall modification involved in abscission (p value = 7.28×10^−04^) and indole-3-acetic acid amido synthetase activity (p value = 1.52×10^−03^) (S5, worksheet 3 cells with yellow background colour). The down-regulated DEG_TS(3S)_ were enriched for terms such as cellular response to iron starvation (p value = 7.03×10^−07^) and iron ion binding (p value = 6.37×10^−04^) (S5, worksheet3 cells with blue background colour). The molecular functions related to metal binding, interestingly had a focus on calcium ion binding in salt absent shoots (S5, worksheet1 cells with yellow background colour) while it is more DNA and Ferric ion binding for salt stressed roots (S5, worksheet2 cells with yellow background colour) and shoots (S5, worksheet3 cells with yellow background colour), respectively.

The transgene dependent salt responsiveness was investigated for the combined effect of both *AtCIPK16* overexpression and salt on the plant. DEG_INT(3R)_ were enriched for response to chitin (p value = 3.08×10^−26^) (S5, worksheet 4 biological process). The pathways the DEG_INT(3R)_ fall in included “carbon metabolism”, “Phenylpropanoid biosynthesis”, “Glyoxylate and dicarboxylate metabolism” and “Galactose metabolism” (S6 column E). Response to carbohydrate stimulus (p value = 5.13×10^−10^) was a GO category identified for the DEG_INT(3S)_ (S5, worksheet 5 biological process). Peroxidases that are involved in the Phenylpropanoid biosynthesis pathway and genes involved in flavonoid biosynthesis were identified through pathway analysis (S6 column F). Furthermore, Calcium-binding EF-hand motif containing genes involved in plant-pathogen interaction were among the pathways which DEG_INT(3S)_ were grouped into (S6, column F).

Next specific roles of DEGs were investigated in the following functional categories; a) transporters/channels, b) regulation of transcription, c) metal handling, d) enzyme families, e) hormone metabolism and f) signalling pathways (S7).

#### 3.5.1 Transporters/Channels

More transporters were identified as DEG_TS(3R)_ compared to transporter DEG_TS(3S)_ and DEG_TC(3S)_ (S7 worksheet 1, under Transport in S8 a, b, d and e). The transporter genes from the DEG_TC(3S)_ included *SLAH3*. The transporter DEG_TS(3R)_ included several *NRTs*, *CHX17*, *CNGC19*, *root hair specific 2* genes. Furthermore, there are transporters from DEG_TS(3R)_ associated with JAZ proteins that are involved in ubiquitination leading to proteolysis, as well as SAUR protein coding genes involved in cell expansion through auxins (S9 a). Other pathways that DEG_TS(3R)_ belonged to were: Phenylpropanoid biosynthesis and phenylalanine metabolism (S10 a) and “valine, leucine and isoleucine degradation related genes”. While transporter DEG_TS(3S)_ are directly or indirectly associated with cell wall biosynthesis, α-Linolenic acid metabolism and Pentose and glucuronate interconversions were associated with DEG_INT(3S)_ (S10 a).

#### 3.5.2 Regulation of Transcription and DNA/DNA Processing

The largest number of DEGs encoding transcription factors (TFs) belonged to DEG_TS(3R)_ (S7 worksheet 2, under regulation of transcription in S8 d). Only five of these TFs were identified in pathways and they fell into plant hormone transduction and plant-pathogen interaction pathways (S10 b). The TFs from DEG_TS(3S)_ were related to limonene and pinene degradation, ubiquitin mediated proteolysis, starch and sucrose metabolism and stilbenoid, diarylheptanoid and gingerol biosynthesis (S10 b). The TF genes from DEG_INT(3R)_ and DEG_INT(3S)_ are directly or indirectly involved in plant-pathogen interactions, starch and sucrose metabolism and plant hormone transduction (S10 b). The hormone signal transduction related genes from DEG_INT(3R)_ are related to auxin, ABA and jasmonic acid (S9 b). RNA synthesis and processing genes were not in DEG_INT(3R)_ while they were in DEG_TS(3R)_ (under RNA synthesis and RNA processing in S8 d and h). DEG_TC(51S)_ contained DNA replication and nucleotide excision repair pathway genes, which included the transcription factor NF-YB11 (*AT2G27470*) (S6 column J).

#### 3.5.3 Metal Synthesis and Assimilation

There are metal related genes within both DEG_TS(3R)_ and DEG_TS(3S)_ which are not in DEG_TC(3R)_ or DEG_TC(3S)_ (under metal handling in S8 a, d and e). Moreover, DEG_INT_ at 3 hours contain metal handling genes that are iron (Fe) related (S7 worksheet 3). Pathways these metal binding DEG_INT_ directly or indirectly modulate include Porphyrin and chlorophyll metabolism (S10 c).

#### 3.5.4 Enzyme Families

There were ‘enzyme related’ 1 DEG_TC(3R)_ and 2 DEG_TC(3S)_ (S7 worksheet 4, under enzyme families in S8 a). However, there are at least 12 ‘enzyme related’ DEG_TS_ (S7 worksheet 4, under ‘enzyme families’ in S8 d and e). Enzyme related DEG_INT(3R)_ were fewer compared to those from DEG_TS(3R)_ but the number of enzyme related genes from DEG_INT(3S)_ and DEG_TS(3S)_ were more or less similar (S7 worksheet 4). ‘Enzyme family’ DEG_TS_ showed associations to genes that fell into pathways of ROS mediation, pathogen interactions and cell growth, and cell wall strengthening (S10 d). Phenylalanine metabolism and phenylpropanoid biosynthesis were seen to be pathways the differentially expressed enzymes of 3 hour DEG_INT_ grouped into (S10 d).

#### 3.5.5 Hormone Metabolism

The hormone related DEG_TC(3S)_ were directly or indirectly involved in ethylene, auxin and brassinosteroid metabolism (S8 j, S9 c). Additionally, 66 of the DEGs from this contrast have putative involvement in biotic stress which include the ethylene signalling related genes and ethylene-responsive element binding protein family genes (S8 q). Within DEG_TS(3R)_ there were genes that were either directly associated to or indirectly modulating genes related to gibberellin, ethylene, auxin, brassinosteroids and JA (S8 l, S9 d). Several genes encode products that are known to be involved in ethylene biosynthesis (*1-aminocyclopropane-1-carboxylic acid (acc) synthase 6; AT4G11280*) and JA biosynthesis (*allene oxide cyclase 2; AT3G25770, allene oxide synthase; AT5G42650*) were evident within DEG_TS(3R)_ (S7 worksheet 5). The potential function of the proteins encoded by these genes mainly was ubiquitination mediated proteolysis (S9 d). However, it was observed that there are genes related to auxin metabolism that may also be related to ubiquitination related proteolysis or plant growth from DEG_TS(3S)_ and DEG_INT(3R)_ (S8 m, o and p; S9 e and f). Plant pathogen interactions were suggestive as a function of the proteins encoded by DEG_TC(3S)_ and DEG_TS(3R)_ (S10 f). A gene of which the product is regulated by ethylene and JA (CEJ1; AT3G50260) was differentially expressed as a DEG_INT(3S)_ (S7 worksheet 5). Furthermore, the only hormone related gene that was differentially expressed in shoots DEG_TS(51S)_ was GASA14 (*AT5G14920*) (S7 worksheet 5, S8 n).

#### 3.5.6 Putative biotic stress related signaling pathways

Compared to the number of signalling related genes in putative biotic stress pathways from DEG_TC(3S)_, there were fewer numbers in DEG_TS_ (S7 worksheet 6, under signalling in S8 q, r and s). Calcium signalling genes dominated the biotic stress related signalling pathway DEG_TC(3S)_ (S7 worksheet 6). Additionally to the groups of genes in putative biotic pathways that were a DEG_TS(3S)_, DEG_TS(3R)_ had genes related to ROS mediation, signal recognition and propagation to the MAPK cascade and heat shock (S8 q, r and s). While DEG_TS(3S)_ were involved in starch and sucrose metabolism, DEG_INT_ contained genes in phenylpropanoid biosynthesis in both roots and shoot (S10 g).

### 3.6 Narrowing Down on Potential Genes Involved in the AtCIPK16 Dependent Salt Response

A pairwise comparison of DEG_INT(3R)_ with the DEG_TC(3R)_ revealed that there are 187 genes out of the 231 genes that are only expressed in a transgene dependent manner in salt (S11 worksheet 1). Furthermore, out of the 152 DEG_INT(3S)_, 120 are uniquely expressed as a transgene dependent salt response, compared to the transgene effect on non-stressed plants (S11 worksheet 2).

The GO terms such as response to ethylene activated signalling pathway, response to wounding and response to chitin were enriched for this subset of 187 genes from root at 3 hours (S11 worksheet 3). Functional clustering of these 187 genes in DAVID revealed the presence of 24 transcription factors, 10 ethylene responsive genes and 15 iron related genes (S11 worksheet 4).

The subset of 120 genes from shoot 3 hours were enriched for GO terms such as cellular response to iron starvation, response to chitin and iron ion homeostasis (S11 worksheet 3). The functional clustering in DAVID revealed that the 120 subset contains genes involved in ‘nucleus’, ‘metal binding’ and ‘transcription regulation’ (S11 worksheet 4).

### 3.7 Narrowing Down on Transcription Factors Putatively Controlled by AtCIPK16

AtCIPK16 was thought to be directly phosphorylating one or more transcription factors in the presence of salinity. To investigate if this was the case, transcription factors with a significant transgene effect only in salt responsiveness were identified; the TFs from DEG_INT_ were compared to TFs from DEG_TC._ Any genes that were common to these two sets were thought to be differentially expressed due to the transgene, yet not explicitly due to transgene effect in salinity. On the other hand, TF genes that were exclusively DEG_INT_ from both roots and shoots at 3 hours were considered as explicitly expressed due to transgene n presence of salt. There were 25 and 16 TFs that were thus, exclusive to DEG_INT(3R) and_ DEG_INT(3S)_, respectively (S7 worksheet 2; yellow background). Interestingly, there was only one such exclusive TF gene common to DEG_INT(3R)_ and DEG_INT(3S)_ (*AT2G25900*; *AtTZF1*) (S7 worksheet 2; yellow background, bold with black border).

It was previously shown that AtTZF1 acts as a transcription factor and binds ARE promoter domains in AU rich regions Pomeranz (Pomeranz et al., 2011; Qu et al., 2014). Therefore, in order to identify potential downstream transcriptional regulatory targets of AtTZF1 in *AtCIPK16* overexpression lines, the region 3000 bp upstream of the transcription start site of all root and shoot transgene dependent salt responsive genes was scanned for the ARE motif through the FIMO tool in MEME suite. In roots 14 such genes with 17 putative ARE promoter motifs were discovered (Table 2). In shoots 10 genes with 13 putative promoter ARE motifs were identified (Table 2).

**Table 2.**
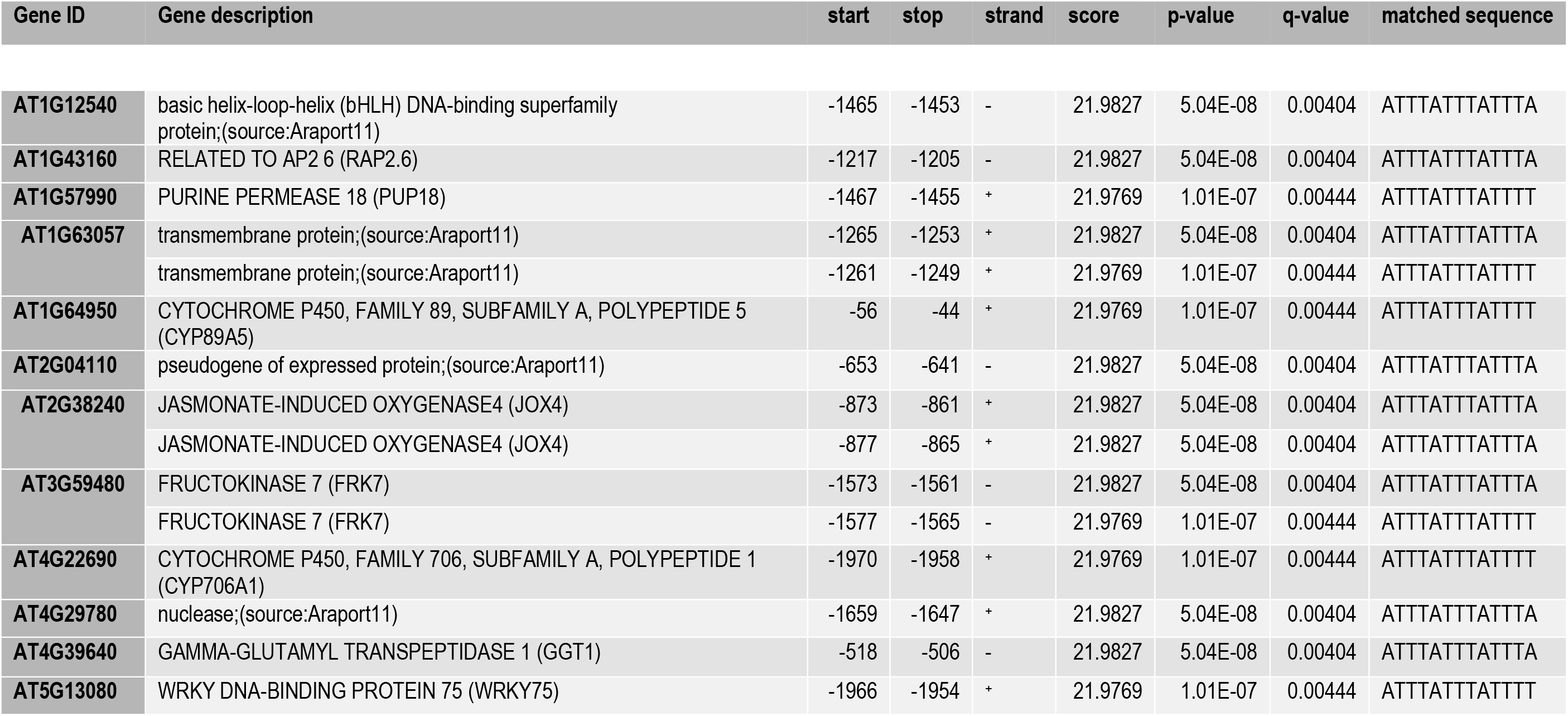
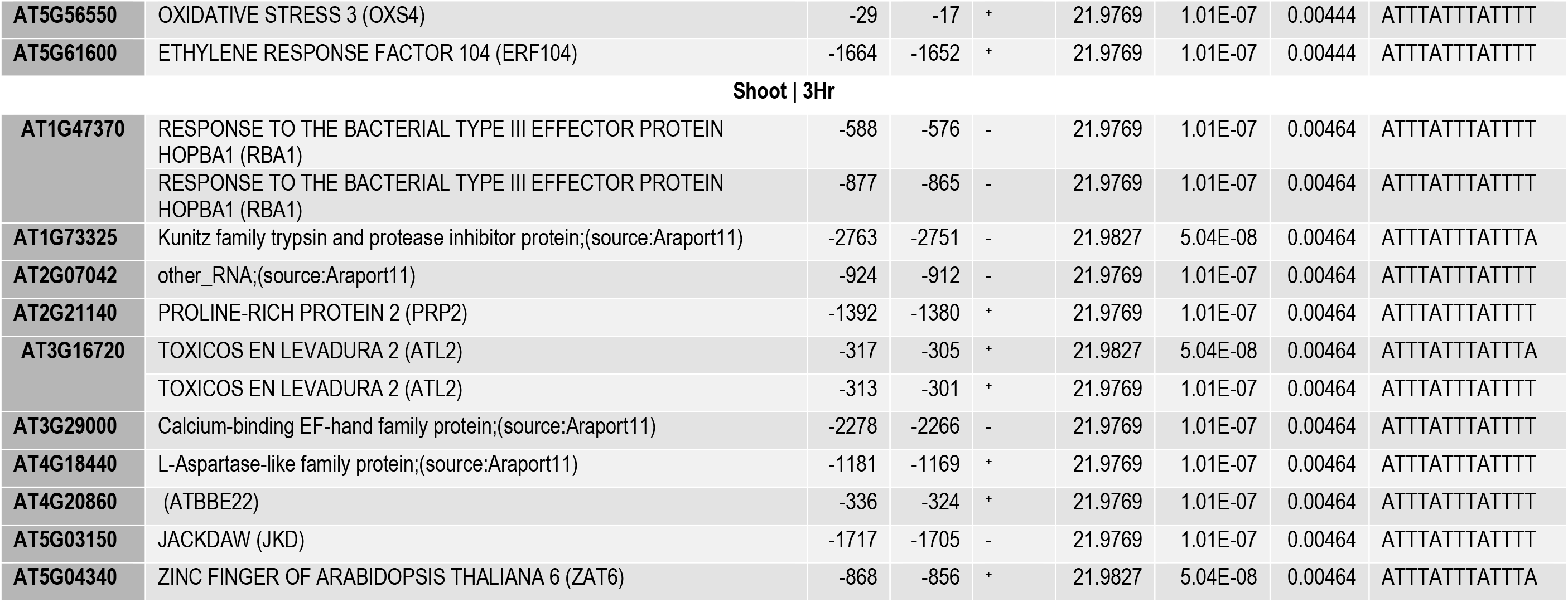
The genes that are putatively regulated by transcriptional activity of AtTZF1 from roots and shoots at 3 hours. The DNA binding motif of AtTZF1 ATTTATTTATTT[T|A] (Pomeranz et al. 2011; Qu et al. 2014), was scanned on the 3,000 bp upstream from the transcription start site (TSS) of all transgene dependent salt responsive genes from roots and shoots at 3 hours (sequences were retrieved using bulk sequence retrieval option from TAIR portal; https://www.arabidopsis.org/tools/bulk/index.jsp and scanned through FIMO from MEME suite; http://meme-suite.org/tools/fimo). The genes with one or more positive hits with a p value ≤ 0.01 are reported here. The gene ID and the descriptions are in the first two columns. The start and the stop site of the identified motif are in the 3^rd^ and 4^th^ columns, respectively. The strand the motif is predicted on is in the 5^th^ columns. FIMO assigned score, p-value and FDR corrected p-value (q-value are in the 6^th^ 7^th^ and 8^th^ columns, respectively. The matched motif is displayed in 9^th^ column.

| Gene ID | Gene description | start | stop | strand | score | p-value | q-value | matched sequence |
| --- | --- | --- | --- | --- | --- | --- | --- | --- |
| AT1G12540 | basic helix-loop-helix (bHLH) DNA-binding superfamily protein;(source:Araport11) | -1465 | -1453 | - | 21.9827 | 5.04E-08 | 0.00404 | ATTTATTTATTTA |
| AT1G43160 | RELATED TO AP2 6 (RAP2.6) | -1217 | -1205 | - | 21.9827 | 5.04E-08 | 0.00404 | ATTTATTTATTTA |
| AT1G57990 | PURINE PERMEASE 18 (PUP18) | -1467 | -1455 | + | 21.9769 | 1.01E-07 | 0.00444 | ATTTATTTATTTT |
| AT1G63057 | transmembrane protein;(source:Araport11) | -1265 | -1253 | + | 21.9827 | 5.04E-08 | 0.00404 | ATTTATTTATTTA |
|  | transmembrane protein;(source:Araport11) | -1261 | -1249 | + | 21.9769 | 1.01E-07 | 0.00444 | ATTTATTTATTTT |
| AT1G64950 | CYTOCHROME P450, FAMILY 89, SUBFAMILY A, POLYPEPTIDE 5 (CYP89A5) | -56 | -44 | + | 21.9769 | 1.01E-07 | 0.00444 | ATTTATTTATTTT |
| AT2G04110 | pseudogene of expressed protein;(source:Araport11) | -653 | -641 | - | 21.9827 | 5.04E-08 | 0.00404 | ATTTATTTATTTA |
| AT2G38240 | JASMONATE-INDUCED OXYGENASE4 (JOX4) | -873 | -861 | + | 21.9827 | 5.04E-08 | 0.00404 | ATTTATTTATTTA |
|  | JASMONATE-INDUCED OXYGENASE4 (JOX4) | -877 | -865 | + | 21.9827 | 5.04E-08 | 0.00404 | ATTTATTTATTTA |
| AT3G59480 | FRUCTOKINASE 7 (FRK7) | -1573 | -1561 | - | 21.9827 | 5.04E-08 | 0.00404 | ATTTATTTATTTA |
|  | FRUCTOKINASE 7 (FRK7) | -1577 | -1565 | - | 21.9769 | 1.01E-07 | 0.00444 | ATTTATTTATTTT |
| AT4G22690 | CYTOCHROME P450, FAMILY 706, SUBFAMILY A, POLYPEPTIDE 1 (CYP706A1) | -1970 | -1958 | + | 21.9769 | 1.01E-07 | 0.00444 | ATTTATTTATTTT |
| AT4G29780 | nuclease;(source:Araport11) | -1659 | -1647 | + | 21.9827 | 5.04E-08 | 0.00404 | ATTTATTTATTTA |
| AT4G39640 | GAMMA-GLUTAMYL TRANSPEPTIDASE 1 (GGT1) | -518 | -506 | - | 21.9827 | 5.04E-08 | 0.00404 | ATTTATTTATTTA |
| AT5G13080 | WRKY DNA-BINDING PROTEIN 75 (WRKY75) | -1966 | -1954 | + | 21.9769 | 1.01E-07 | 0.00444 | ATTTATTTATTTT |
| **AT5G56550** | OXIDATIVE STRESS 3 (OXS4) | -29 | -17 | + | 21.9769 | 1.01E-07 | 0.00444 | ATTTATTTATTTT |
| **AT5G61600** | ETHYLENE RESPONSE FACTOR 104 (ERF104) | -1664 | -1652 | + | 21.9769 | 1.01E-07 | 0.00444 | ATTTATTTATTTT |
| **Shoot \| 3Hr** |  |  |  |  |  |  |  |  |
| **AT1G47370** | RESPONSE TO THE BACTERIAL TYPE III EFFECTOR PROTEIN HOPBA1 (RBA1) | -588 | -576 | - | 21.9769 | 1.01E-07 | 0.00464 | ATTTATTTATTTT |
|  | RESPONSE TO THE BACTERIAL TYPE III EFFECTOR PROTEIN HOPBA1 (RBA1) | -877 | -865 | - | 21.9769 | 1.01E-07 | 0.00464 | ATTTATTTATTTT |
| **AT1G73325** | Kunitz family trypsin and protease inhibitor protein;(source:Araport11) | -2763 | -2751 | - | 21.9827 | 5.04E-08 | 0.00464 | ATTTATTTATTTA |
| **AT2G07042** | other_RNA;(source:Araport11) | -924 | -912 | - | 21.9769 | 1.01E-07 | 0.00464 | ATTTATTTATTTT |
| **AT2G21140** | PROLINE-RICH PROTEIN 2 (PRP2) | -1392 | -1380 | + | 21.9769 | 1.01E-07 | 0.00464 | ATTTATTTATTTT |
| **AT3G16720** | TOXICOS EN LEVADURA 2 (ATL2) | -317 | -305 | + | 21.9827 | 5.04E-08 | 0.00464 | ATTTATTTATTTA |
|  | TOXICOS EN LEVADURA 2 (ATL2) | -313 | -301 | + | 21.9769 | 1.01E-07 | 0.00464 | ATTTATTTATTTT |
| **AT3G29000** | Calcium-binding EF-hand family protein;(source:Araport11) | -2278 | -2266 | - | 21.9769 | 1.01E-07 | 0.00464 | ATTTATTTATTTT |
| **AT4G18440** | L-Aspartase-like family protein;(source:Araport11) | -1181 | -1169 | + | 21.9769 | 1.01E-07 | 0.00464 | ATTTATTTATTTT |
| **AT4G20860** | (ATBBE22) | -336 | -324 | + | 21.9769 | 1.01E-07 | 0.00464 | ATTTATTTATTTT |
| **AT5G03150** | JACKDAW (JKD) | -1717 | -1705 | - | 21.9769 | 1.01E-07 | 0.00464 | ATTTATTTATTTT |
| **AT5G04340** | ZINC FINGER OF ARABIDOPSIS THALIANA 6 (ZAT6) | -868 | -856 | + | 21.9827 | 5.04E-08 | 0.00464 | ATTTATTTATTTA |

### 3.8 Known DEGs with Potential Phosphorylation Ability with a Focus on MAPK Phosphorylated DEGs

Furthermore, the NetPhoS4.1 phosphorylation prediction server results showed that the above subset of 187 genes from DEG_INT(3R)_ contained 181 genes that code for amino acid sequences containing multiple serine/threonine phosphorylation sites (S11 worksheet 5). Furthermore, NetPhoS4.1 server shows that, 109 out of 120 DEG_INT(3S)_ could potentially be phosphorylated with a given score ≥ 0.9 (S11 worksheet 6). This observation was not surprising because the consensus sequence of a phosphorylation site is less than 20 amino acids long.

The ability of protein phosphorylation is best studied for the MAPK cascade in various stress conditions. Therefore, genes that are phosphorylated by various MPKs were identified. There were twelve and two DEG_INT(3R)_ and DEG_INT(3S)_, respectively, that are potential targets of the MAPK phosphorylation (S12). Majority of the identified substrates are phosphorylated by MPK6 (S12).

While the nine DEGs that are common between DEG_INT(3R)_ and DEG_INT(3S)_ are showing the ability to get phosphorylated (S11 worksheet 5 and 6), ZAT10 (AT1G27730) and ATCTH (AT2G25900) are also known to be substrates of the MAPK cascade (S12).

### 3.9 Co-expression Analysis

#### 3.9.1 Roots

The WGCNA network analysis created 66 modules. Hub genes of a module are comprehended as the key drivers of that module which have highest connectivity to the module (i.e. most responsible for the intact network topology). In order to identify the effect of transgene dependent salt responsiveness on these modules (i.e. gene clusters), hub genes from each cluster were screened for DEG_INT(3R)_. Out of the 86, 14 modules contained one or more DEG_INT(3R)_ as hub genes (S13 worksheet 1) and were selected for further investigations. The genes in each selected module are in S13 (worksheet 2). Hub genes from the modules are shown in S13 (worksheet 3) and the DEG_INT(3R)_ are highlighted in yellow. Since there were no transgene dependent salt responsive genes in roots, no such analysis was performed for the 51 hour time-point.

To extend the network analysis further and retrieve biological relevance underlying the identified modules from 3 hours, functional enrichment analysis of genes in the selected 14 modules was performed (S13 worksheet 4, 5 and 6). The green module that contained 1026 genes was highly enriched for the biological process (BP) response to chitin (p value = 2.87×10^−18^) (S13 worksheet 4). The darkgrey module was enriched for the term ‘response to wounding’ (p value = 3.81×10^−05^) while pink module was enriched for ‘defence response’ (p value = 2.75×10^−09^) (S13 worksheet 4). Interestingly, the lightsteelblue1 module was highly enriched for photosynthesis (p value = 1.89×10^−61^). The other modules were enriched for the terms ‘response to water deprivation’, ‘response to abscisic acid’, ‘response to absence of light’, ‘circadian rhythm’, ‘autophagy’, ‘rRNA modification’, ‘cell wall organisation’, ‘response to karrikin’, ‘oxidation reduction process’ and ‘syncytium formation’ (S13 worksheet 4).

*AtCIPK16 was* found in the yellow module and co-clustered with *AtHKT1* (*AT4G10310)* (S13 worksheet 2) and *trehalose phosphate synthase 10* (*AT1G60140*). The yellow module is enriched for ‘carbohydrate metabolic process’ (p value = 0.002) and ‘sodium ion transport’ (p value = 0.002) (S13 worksheet 4). The KEGG pathways the yellow module genes fall into include starch and sucrose metabolism (S13 worksheet 5).

#### 3.9.2 Shoots

There were 17 WGCNA modules for shoots. Out of these four modules contained transgene dependent salt responsive genes from shoot at 3 hours as hub genes (S13 worksheet 1). Again the analysis was restricted to the 3hr time-point since there were no transgene dependent salt responsive genes in shoots at 51 hours. The module genes and the respective module hub genes that were transgene dependent salt responsive are in S13 (worksheet 8 and 9, respectively).

The tan module was highest enriched for the term cellular response to iron starvation (S13 worksheet 10) and contained *bHLH43* (*POPEYE/PYE: AT3G47640*). The blue module, which also contained *AtCIPK16*, on the other hand was enriched for the term mRNA processing (p value = 4.81×10^−19^) (S13 worksheet 10). Turquoise module was enriched for ribosome biogenesis (p value = 2.33×10^−10^) while magenta was enriched for water deprivation (p value = 2.00×10^−13^) (S13 worksheet 10).

## 4 Discussion

Plant transformation has the potential to be a fast, versatile method to improve plant traits with the ultimate goal of increasing or stabilising crop yield under adverse environmental conditions (Gilliham et al. 2017). It has been shown that *AtCIPK16* overexpression in Arabidopsis conferred enhanced salt tolerance (Roy et al. 2013). However, the underlying molecular mechanisms that govern the observed traits were unknown. It is important to identify the targets which are affected by AtCIPK16, to determine whether overexpression of *AtCIPK16* is not detrimental, but only beneficial to the plant in the long term. We attempted to reduce this disparity in knowledge using a transcriptomic approach.

The experiment was designed to study the transcriptome differences between the transgenics and null transgenics at two different time points that have possible early (3 hours after initial salt application) and late (51 hours after initial salt application) responses to salinity stress. Illumina sequencing was used to generate the transcriptomic data which were subsequently mapped and analysed to gauge the salt tolerance responses of *AtCIPK16* overexpression.

### 4.1 Effect of *AtCIPK16* transgene in salt stress

We are now able to provide laboratory and *in-silico* evidence to support the assumption - AtCIPK16 may elicit its function within the root cell nucleus in the presence of salt stress and this function includes the manipulation of one or more transcription factors (TFs) as follows: a) we previously showed that AtCIPK16 possesses a putative nuclear localisation signal (Amarasinghe et al. 2016); here we show that b) AtCIPK16 is localised partially to the nucleus; c) AtCIPK16 has a putative DNA binding domain which may bind it to a DNA bound molecule; d) there is minimal gene expression differences in control roots which increases almost 4 fold in salt stressed roots; and, e) a large number of TFs are differentially expressed.

It is likely that a regulator is needed to release the AtCIPK16 from its auto-inhibitory status and direct towards the targets; however, RNA-Seq experiments cannot identify the potential regulators of AtCIPK16. Nonetheless, especially in roots it could possibly be that, these regulators are dormant until the plant is stressed. Lee et al. (2007) has suggested the possibility of CBL1 and CBL9 to be the interacting partners of CIPK16. More recently, the ability of other kinases, such as GRIK kinases, to release the auto-inhibitory state of SOS2 has been established (Barajas-Lopez et al. 2018). This implies that there could be an alternative interactome for CIPKs apart from the well-known CBLs to release it from its’ auto-inhibitory form.

### 4.2 Possible Downstream Activation of AtTZF1

Among the TFs differentially expressed, we identified one CCCH zinc finger (*AtTZF1*) that stands out as being the only upregulated TF in both roots and shoots at 3 hours exclusive to the transgene dependent salt responsiveness. A previous study revealed that Arabidopsis plants overexpressing At*TZF1* showed enhanced salinity tolerance compared to the wild type due to reduced shoot Na^+^ accumulation, increased chlorophyll content and increased growth (Han et al. 2014). *AtCIPK16* transgenics also did show reduced Na^+^ accumulation and increased biomass (Roy et al. 2013). We propose AtTZF1 as a potential downstream master regulator of AtCIPK16 mediated salt stress tolerance and suggest knockout or knockdown lines to investigate this contention. Activation of C3H zinc fingers by post translational phosphorylaion has been shown and suggested previously for plants and mammals (Bogamuwa and Jang 2016; Brooks and Blackshear 2013; Maldonado-Bonilla et al. 2014; Taylor et al. 1995). It was identified that a serine downstream of the zinc finger can be phosphorylated and enhance the activity of the TF (Cziferszky et al. 2002). It would be interesting to know whether AtTZF1 enhances its activity through phosphorylation, and if so, could AtCIPK16 phosphorylate AtTZF1 as well. Furthermore, in this study it was shown that there are 14 and 10 genes from roots and shoots respectively that could be transcriptionally regulated by AtTZF1 in presence of AtCIPK16. This is an exciting path for further investigations because manipulation of a fine-tuned TF that can control many downstream targets is a desirable feature for the development of stress-tolerant crops without detrimental consequences or yield penalty (Zhou et al. 2007).

### 4.3 Potential Regulation through Phytohormones

It was evident from the functional categorisation that hormone metabolism related genes, mainly those related to ethylene biosynthesis (e.g. 1-Aminocyclopropane-1-carboxylic acid synthase 6; ACC synthase 6/ACS6) (Wang et al. 2002), jasmonic acid (JA) biosynthesis and cross talk with ethylene (e.g. Allene Oxide Synthase/AOS, AtERF1, CEJ1 and AtMYC2) [79–82] and auxin regulation (e.g. SAUR genes) (Ren and Gray 2015), were differentially expressed in the transgenic salt stressed transcriptome, especially at 3 hours. It could mean that phytohormone regulation is an important aspect of AtCIPK16 mediated salt stress tolerance. Auxin responsive factors such as SAUR proteins are well established to be involved in cross-talk between biotic and abiotic stress tolerance (Bouzroud et al. 2018; Ghanashyam and Jain 2009; Jain and Khurana 2009; Saini et al. 2017). There is also a possibility that while AtCIPK16 affects the transcription of these genes downstream, the phosphorylation also could mediate their activity post-translationally. Salt stress was shown to increase ethylene production (Cao et al. 2007; Cao et al. 2006). In turn, ethylene biosynthesis and signalling has been shown to reduce salt sensitivity (Cela et al. 2011; Tao et al. 2015). Could it be that the downstream activity of AtCIPK16 under salt stress promotes the ethylene biosynthesis at least partly owing to increased ACS6 gene expression? If so, higher accumulation rates of ethylene may inhibit the negative effect of ethylene receptors on salinity caused growth arrest. *AtCIPK16* overexpression in *ACS6 knockouts* can firmly link the function of ACS6 to salinity tolerant *AtCIPK16* overexpressing phenotypes.

Inhibition of primary root growth and promotion of lateral root growth owing to redistributed auxin from shoots to roots as a response to ethylene synthesis has been suggested to be important in tolerating low salinity stress in Arabidopsis (Zhao et al. 2011) and more recently in barley (Witzel et al. 2018). The involvement of a protein encoded by another *CIPK* gene (*SOS2*; *AtCIPK24*) in the process of auxin redistribution that contributes to lateral growth development in mild salinity stress has been reported previously (Zhao et al. 2011). Data on lateral root length and number could provide answer to the query on whether AtCIPK16 mediated salt tolerance cause a similar morphological change.

A zinc finger protein named ZFP5 has been shown to integrate ethylene with other phytohormones to control root hair development in Arabidopsis (An et al. 2012). In the present study, ZFP5 is co-expressed and a direct neighbour of AtZAT10 in the green module of roots. It is possible that AtZAT10 may be involved in modulating the activity of ZFP5. If so, the downstream effect of *AtCIPK16* overexpression may direct the ethylene signalling cascade towards ZFP5 through AtZAT10 which can cause morphological effects such as root hair growth. Plant root hair growth is observed in salinity stress which enables rapid influx and efflux of ions (An et al. 2012; Gilroy and Jones 2000). It has been suggested that root hairs show preferential expression of certain K^+^ channels involved in K^+^ uptake (Ivashikina et al. 2001). Increase of K^+^ increases the K^+^/Na^+^ ratio hence provides a protective function against the toxicity of Na^+^ within the cytosol (Maathuis and Amtmann 1999). This could be investigated further in transgenic lines by measurements of K^+^ in the roots in control and salt stressed conditions.

### 4.4 Transgenics Adapt to New Conditions Faster than the Wild type

There could be several possible reasons for not seeing any DEG_INT_ at 51 hours. Firstly, *AtCIPK16* mediated salinity tolerance could have already reached homeostasis by 51 hours, while it still has not reached homeostasis in the null transgenics; this could explain why we saw reduced number of DEGs as an effect of salt in transgenics compared to the nulls at 51 hours, in contrast to 3 hours. Rapid adjustment to new conditions may explain the high salinity tolerance of halophytes, such as mangroves, and this may be an important mechanism for improved salt tolerance (Krishnamurthy et al. 2017; Liang et al. 2012; Zhu 2001). Secondly, the rapid onset of the initial osmotic adjustments to counteract the immediate reduction in plant growth due to salinity stress, requires instant root-to-shoot signalling once salt has been detected at the roots (Batistič and Kudla 2010; Gilroy et al. 2014; Roy et al. 2014; Roy et al. 2011; Shabala et al. 2016). It is plausible that overexpression of *AtCIPK16* may be responsible for the prompt induction of its downstream stress tolerant pathway that aides in the rapid adjustment to the new stressful condition.

### 4.5 Potential of *AtCIPK16* to be an Conciliator between Abiotic and Biotic Stress Responses

In nature, plants are exposed to various concurrent abiotic and biotic stresses. Abiotic stresses have been shown to affect the tolerance of biotic stresses negatively as well as positively. Cross-tolerance is a term used to define the phenomenon of abiotic stress augmenting plant pathogen resistance (Ayres 1984). An example for this is barley, *Hordeum vulgare*, grown in saline water exhibiting enhanced tolerance to the barley powdery mildew fungus (Wiese et al. 2004) while, pre-treatment of Arabidopsis with chitin, a key component of the fungal cell wall, was shown to improve salt tolerance (Brotman et al. 2012). Recently, the identified interaction of a chitin receptor CERK1 with the Na^+^ induced Ca^2+^ channel ANN1, was shown to function both in fungal attack and salt stress tolerance (Espinoza et al. 2017). Seeing transgene dependent salt responsive genes involved in putative biotic stress pathways poses the question whether the *AtCIPK16* mediated molecular mechanism also confers tolerance to biotic stresses. This is plausible due to the fact that *AtCIPK16* overexpression, leads to an increased abundance of the respective CIP kinase which can phosphorylate multiple targets, hence could potentially activate more than one pathway or signal transduction cascade and lead to cross-tolerance. Transgene dependent salt responsive genes being enriched for chitin response in both roots and shoots would support our speculation on *AtCIPK16’s* overexpression also activating biotic stress tolerance pathways. There is previous evidence and suggestions on the involvement of CIPKs such as CIPK11 (Xie et al. 2010), CIPK25 (Huibers et al. 2009) and CIPK26 (Drerup et al. 2013) in biotic stresses.

We found an abundance of DEGs implicated in the phenylpropanoid biosynthesis pathway in salinity stressed roots and shoots. Phenylpropanoids are considered antimicrobial compounds that were shown to increase resistance to viral and bacterial infections or function as signalling molecules in biotic stress responses (Naoumkina et al. 2010). Phenylpropanoid biosynthesis, however, can lead to lignin formation which increases the rigidity of plant cell wall and stalls the plants development (Gall et al. 2015) thereby reducing its biomass. Plants with *AtCIPK16* overexpression however had higher biomass than that wild type plants grown under salinity stress (Roy et al. 2013). Therefore, it is still unclear whether the phenylpropanoid biosynthesis is detrimental or favourable in this particular situation.

### 4.6 The Proposed Model of Salinity Response in *AtCIPK16* Transgenics

We propose a molecular pathway of *AtCIPK16* mediated salt stress tolerance (Figure 8). Salt stress signals may be sensed by “sensor molecules” owing to the salt stress related changes in cytosolic Ca^2+^ levels in root cells. These sensor molecules then interact with/phosphorylate AtCIPK16 to release it from the auto-inhibitory state (Barajas-Lopez et al. 2018; Sanchez-Barrena et al. 2007). The active form of AtCIPK16 would subsequently phosphorylate multiple downstream targets including ACS6 which in return would induce the ethylene biosynthesis. Elevated levels of ethylene then overrule ethylene receptor induced arrest of root growth. Furthermore, ethylene can promote auxin redistribution to promote lateral root and root hair growth which may involve ZAT10/12 and AtZFP5 (An et al. 2012; Ivanchenko et al. 2008; Zhao et al. 2011). A possible increase in root surface area could result in elevated uptake of K^+^ thereby creating a favourable K^+^/Na^+^ ratio (Cellier et al. 2004). Furthermore, carbohydrates such as trehalose may be synthesised in the roots, possibly as osmoprotective molecules. AtCIPK16 may even enhance the activity of AtTZF1 through phosphorylation which leads to regulation of multiple downstream targets of AtTZF1. Through increasing the expression of downstream targets such as *ERF104* in roots, AtTZF1 might aid more ethylene production. Additionally, AtCIPK16 may phosphorylate and mediate the activity of *AOC* which leads to reinforce biosynthesis of JA that can elicit salinity tolerance responses such as inhibition of primary root growth, as well as potential root-shoot signalling. Salt stress signalling to shoots could activate Fe accumulation and suppress the inhibition of photosynthetic systems which in return may promote plant shoot growth. Auxin in shoots may also promote cell growth that increases biomass of AtCIPK16 transgenics in salinity.

**Figure 8.**
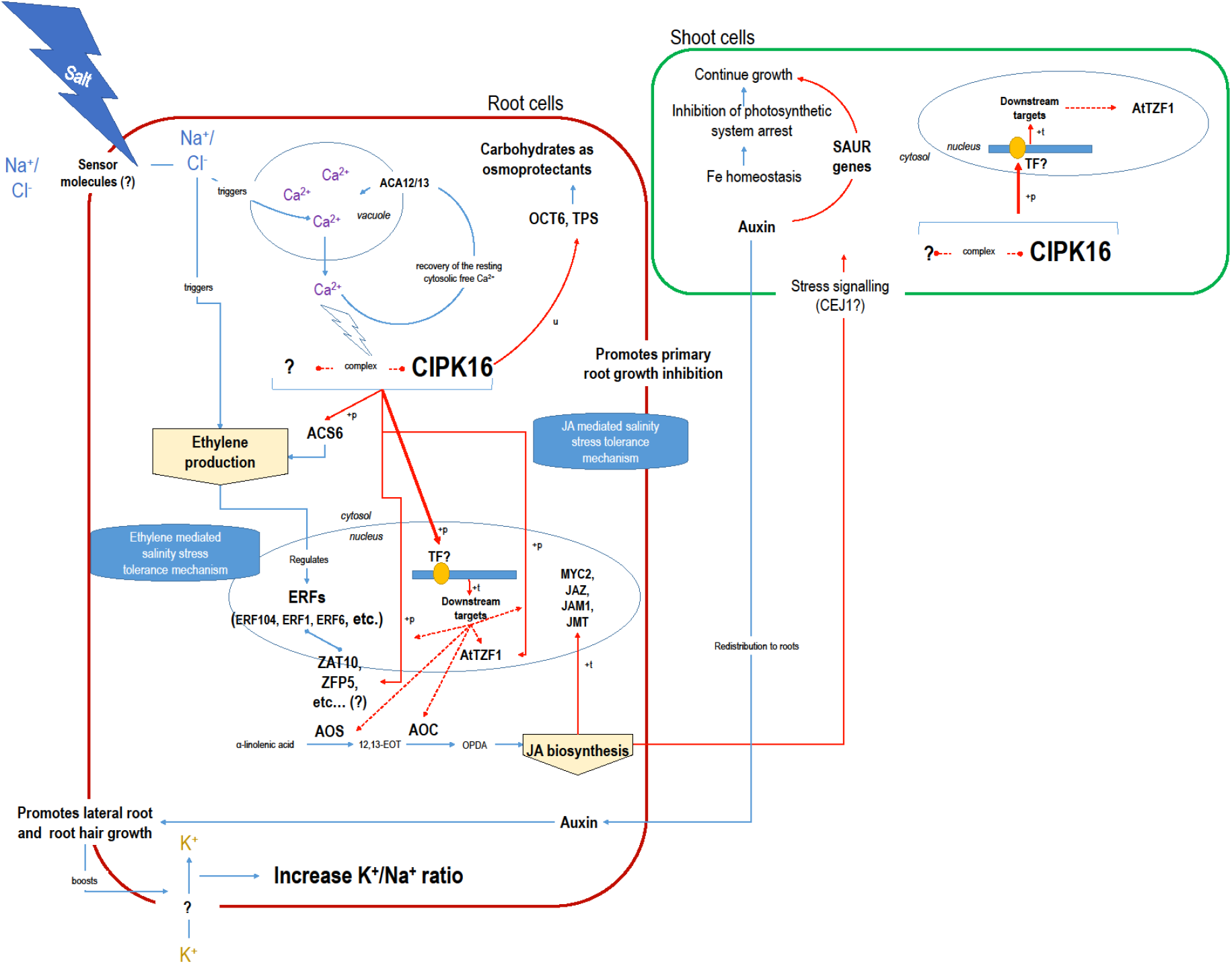
The proposed model for *AtCIPK16* mediated salinity tolerance mechanism in Arabidopsis. The model proposes the involvement of ethylene and JA in the *AtCIPK16* overexpression mediated salinity tolerance. Blue arrows depict currently known knowledge and red arrows depict the proposed AtCIPK16 related pathways. If a potential method of regulation is known for the proposed pathways they are shown next to the arrow (^+^p: phosphorylated for enhanced activity; ^+^t: enhanced by transcriptional regulation; u: unknown method of regulation). The arrow heads represent the direction of regulation. Double pointed arrows are when the direction of regulation is uncertain.

### 4.7 Conclusion

We now have reasons to believe that *AtCIPK16* mediated salinity tolerance is achieved through the activity of a host of TFs synchronised with the regulation of phytohormones, mainly ethylene and jasmonic acid. Modulating TFs and phytohormone mediated responses may well be a crucial aspect in generating salt tolerant germplasm. However, this can be an audacious task, given that both these components are involved in all aspects of a plant’s life cycle including its response towards environmental cues. Yet, their importance is re-established by this study. The large overlap of putative functionality of differentially expressed gene products with biotic stress responses shows that *AtCIPK16* overexpression may have the ability to elicit tolerance to multiple abiotic and biotic stresses which is also an important trait towards developing a field-ready salt tolerant plant. However, the importance of *AtCIPK16* as a genetic tool for engineering salt tolerance in crops such as barley needs further investigation. These investigations should shed light on the stability of the transgene in propagation through generations, its ability to be fine-tuned by using cell specific promoters and thereby eliminating any negative effects of *AtCIPK16* overexpression.

## Acknowledgements

This project was funded by the Grains Research and Development Corporation (GRDC): project UA000145 to SJR and MG. This research was also supported by the Australian Research Council (ARC) and GRDC funding to the Australian Centre for Plant Functional Genomics (ACPFG), and ARC projects IH130200027 to UB and SJR, and CE1400008 to MG. We further thank ACPFG and The University of Adelaide for providing the PhD scholarship to SLA and The University of Adelaide for providing the Adelaide Graduate Scholarship to WH. Authors also wish to thank Mr. Ashan Hettiarachchige for creating the final high resolution images according to the guidelines from the journal.

## Supplementary Material

**S1: Summary of *k-mer* baiting step to confirm the presence of the transgene in samples** a) Construct architecture of the *AtCIPK16* transgene; b) the sequence in between the *35S* promoter and the *AtCIPK16* exon 1; c) *35S* promoter sequence that was used to bait the 5^′^ UTR region of the transgene; d) *NOS terminator* sequence that was used to bait the 3^′^ UTR region of the transgene; e) wild type *AtCIPK16* 5^′^ UTR region; f) wild type *AtCIPK16* 3^′^ UTR region; g) baiting results from the 3 hour time point; h) baiting results from the 51 hour time point; for g and h the columns from left to right represent the following: *column1*: name of the fastq file, *column2*: number of baits for the transgene 3^′^ UTR region, *column3*: number of baits for the transgene 5^′^ UTR region, *column4*: number of baits for the transgene 5^′^ UTR region between exon1 and the *35S* promoter, *column5*: number of baits for the wild type 3^′^ UTR region, *column6*: number of baits for the transgene 5^′^ UTR region, *column7-9*: experimental conditions of each sample. Rows of g and h are coloured for green shades to represent the shoot samples and brown shades to represent root samples.

**S2: Percentages of mapped, multi-mapped and unmapped reads from the samples using STAR aligner**

**S3: The plot to show the justification of the removal of a root and a shoot sample**

a) Mapped raw counts for *AtCIPK16* (*AT2G25090*) of the 3 hour samples, b) Normalised counts mapped to *AtCIPK16.* The blue semi-transparent bars indicate the samples that were removed based on their visibly high amount of normalised reads mapped to *AtCIPK16*.

**S4: Differentially expressed genes (DEGs) resulted from the hypothesis testing shown in Figure 5**

Where applicable the yellow colour represents up regulated genes and blue colour represents down regulated genes.

**S5: GO enrichment analysis of the DEGs performed through PlantGSEA online web server (http://structuralbiology.cau.edu.cn/PlantGSEA/index.php)**

Where applicable the yellow colour represents GO terms enriched by up regulated genes and blue colour represents GO terms enriched by down regulated genes.

**S6: Results of Kyoto Encyclopedia of Genes and Genomes (KEGG) information mining for the DEGs using the “Search and Color Pathway” option (http://www.kegg.jp/)**

**S7: Selected MapMan categories that the DEGs fall into**

**S8: MapMan pathway analysis of the DEGs**

a-i: cell function overview; j-p: regulation overview; q-u: putative biotic stress pathways.

**S9: KEGG Pathways of DEG subsets that are discussed in Mapman categories and their associated genes identified through ATTED-II.**

The pathways are auto generated through the search and colour pathway option in Kyoto Encyclopaedia of Genes and Genomes (KEGG) server (www.genome.jp/kegg/) The input genes for each section are the up regulated DEGs for a), c), d) and e). and all DEGs for b) and f). The associated genes that are included in the list are retrieved through the NetworkDrawer tool with default options (Platform is automatic, CoEx option add many genes and PPI option add a few genes) of ATTED-II server (atted.jp/). Rectangular boxes with green colour background represent genes in the pathway, arrows represent a molecular interaction or a relationship. The red framed green boxes with red letters show the genes that are in the input lists, each group of DEGs used to mine the pathways in each sub figure are mentioned below the respective sub figure a)-f), empty circles represent chemical molecules, rectangles with rounded edges shows the ink from the current pathway to another pathway, doubled lines represent the plasma membrane, the dashed grey lines are shown when a direct association between two molecules are unknown.

**S10: KEGG pathways enriched for the DEG subsets of selected MapMan categories and their associated genes identified through ATTED-II**

**S11: Novel genes involved in the AtCIPK16 dependent salt responsiveness**

The gene lists identified from both roots and shoots, their GO enrichment and functional categorisation through DAVID online web server and the novel genes that have the ability to get phosphorylated that are identified through NetPhos3.1b online server.

**S12: Identified novel AtCIPK16 transgene dependent salt responsive genes that are also putative MAPK substrates**

**S13: Summary of the WGCNA analysis of DEGs**

Interesting modules were selected if one or more transgene dependent salt responsive DEGs from the respective tissue (i.e. root or shoot) are hub genes of the said module. The GO enrichment was performed for each selected module through DAVID online tool (https://david.ncifcrf.gov/). The pathway analysis was cone using search and Color pathway option in Kyoto Encyclopedia of Genes and Genomes (KEGG) (www.genome.jp/kegg/).

