## Supplementary material for "AtCIPK16 Mediates Salt Stress Through Phytohormones and Transcription Factors": S1

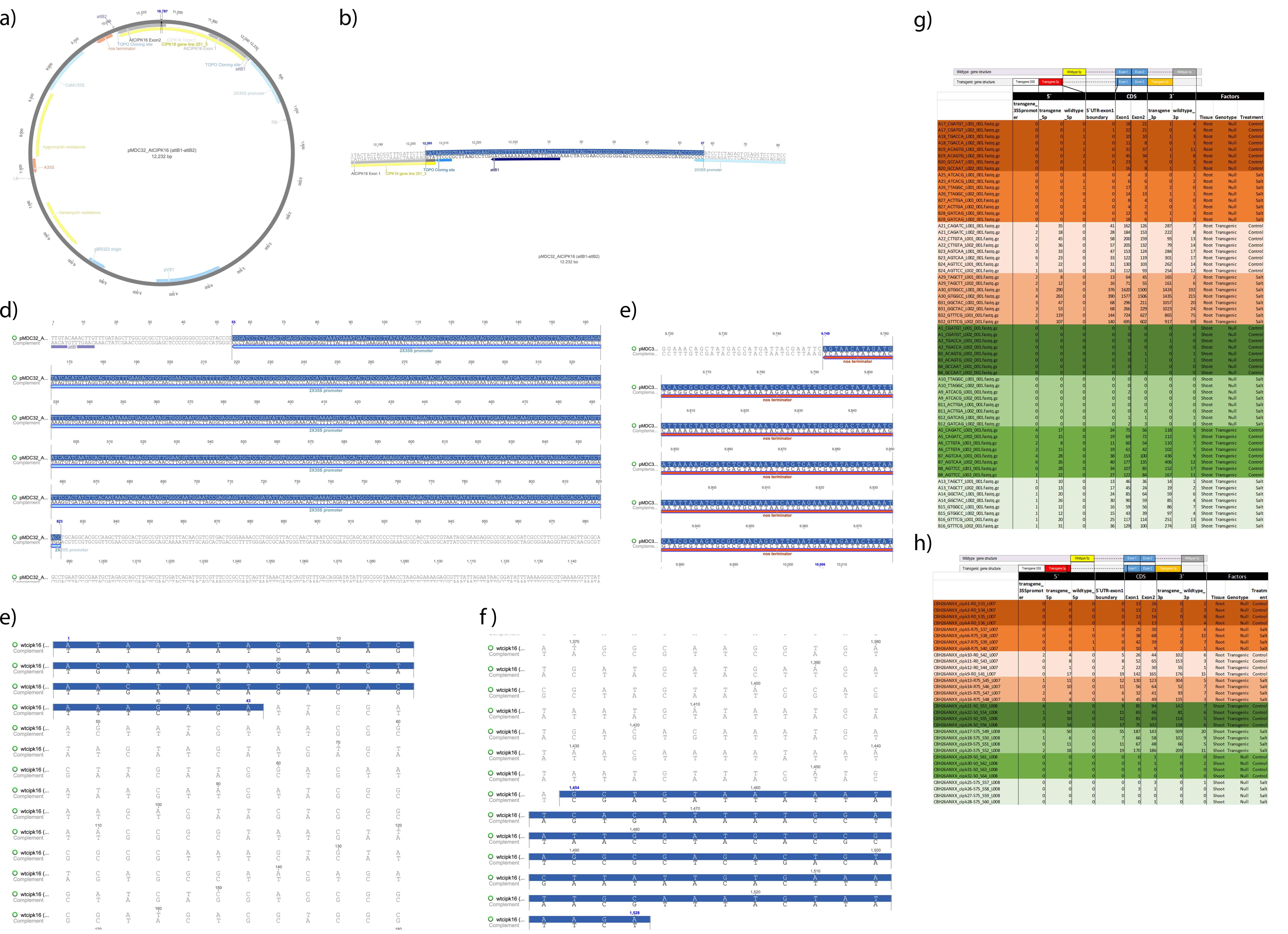


**S1: Summary of *k-mer* baiting step to confirm the presence of the transgene in samples**

a) Construct architecture of the *AtCIPK16* transgene; b) the sequence in between the 35S promoter and the AtCIPK16 exon 1; c) 35S promoter sequence that was used to bait the 5` UTR region of the transgene; d) NOS terminator sequence that was used to bait the 3` UTR region of the transgene; e) wild type *AtCIPK16* 5` UTR region; f) wild type AtCIPK16 3` UTR region; g) baiting results from the 3 hour time point; h) baiting results from the 51 hour time point; for g and h the columns from left to right represent the following: column1: name of the fastq file, column2: number of baits for the transgene 3` UTR region, column3: number of baits for the transgene 5` UTR region, column4: number of baits for the transgene 5` UTR region between exon1 and the 35S promoter, column5: number of baits for the wild type 3` UTR region, column6: number of baits for the transgene 5` UTR region, column7-9: experimental conditions of each sample. Rows of g and h are coloured for green shades to represent the shoot samples and brown shades to represent root samples.
