## Supplementary material for "AtCIPK16 Mediates Salt Stress Through Phytohormones and Transcription Factors": S3

a)

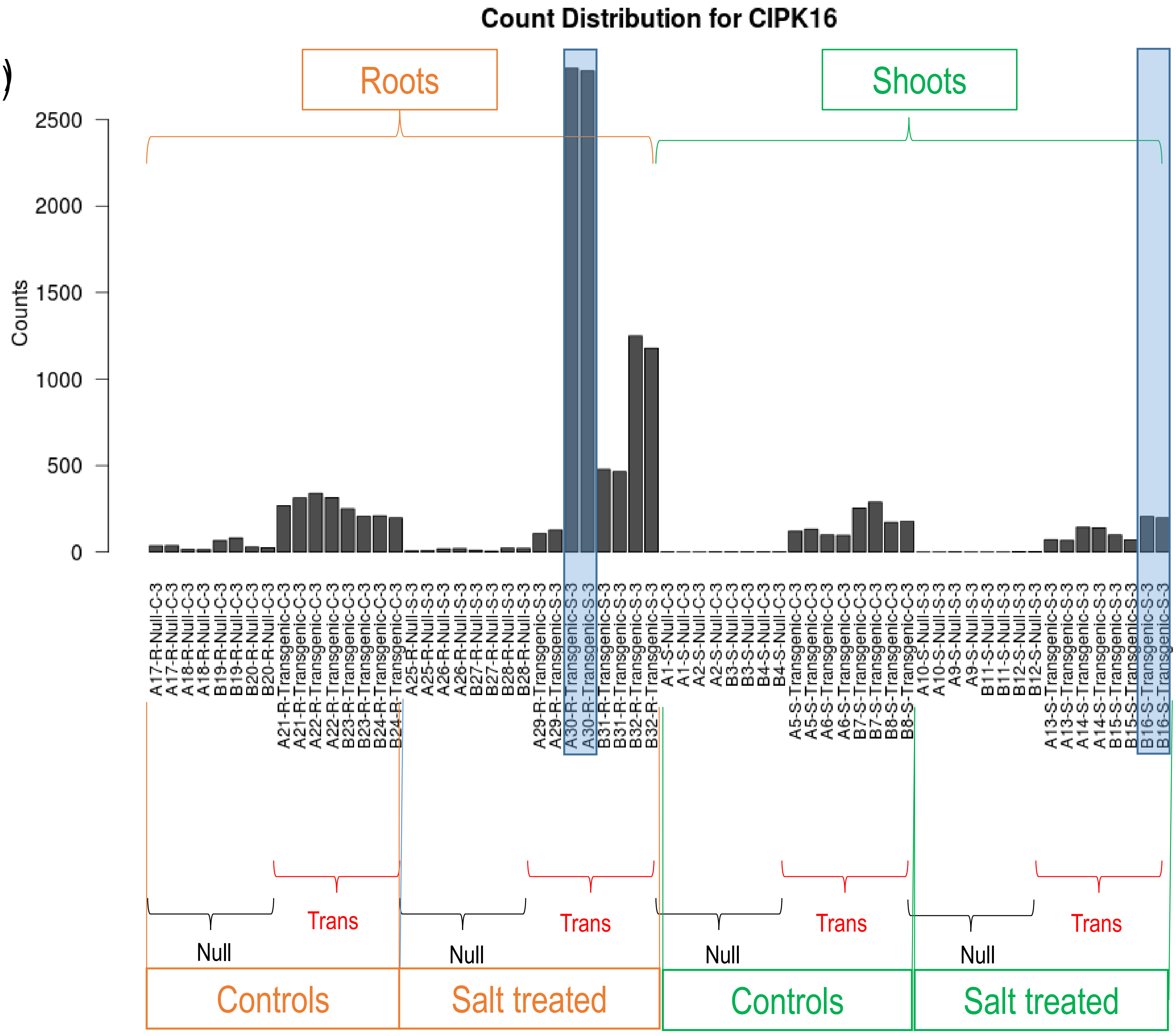


b)

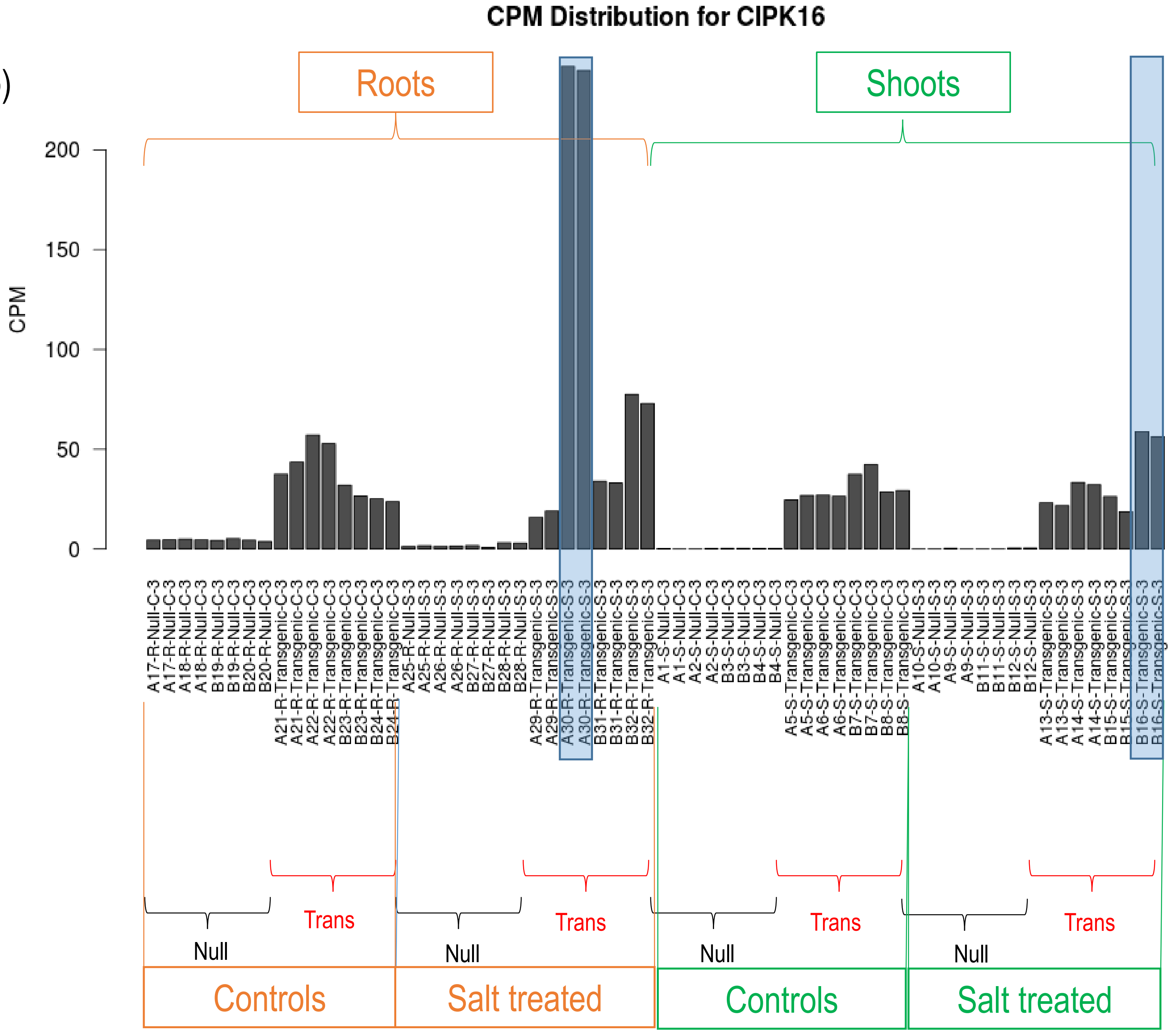


**S3: The plot to show the justification of the removal of a root and a shoot sample**

a) Mapped raw counts for *AtCIPK16* (*AT2G25090*) of the 3 hour samples, b) Normalised counts mapped to *AtCIPK16*. The blue semi-transparent bars indicate the samples that were removed based on their visibly high amount of normalised reads mapped to *AtCIPK16*.
