## Supplementary material for "AtCIPK16 Mediates Salt Stress Through Phytohormones and Transcription Factors": S8

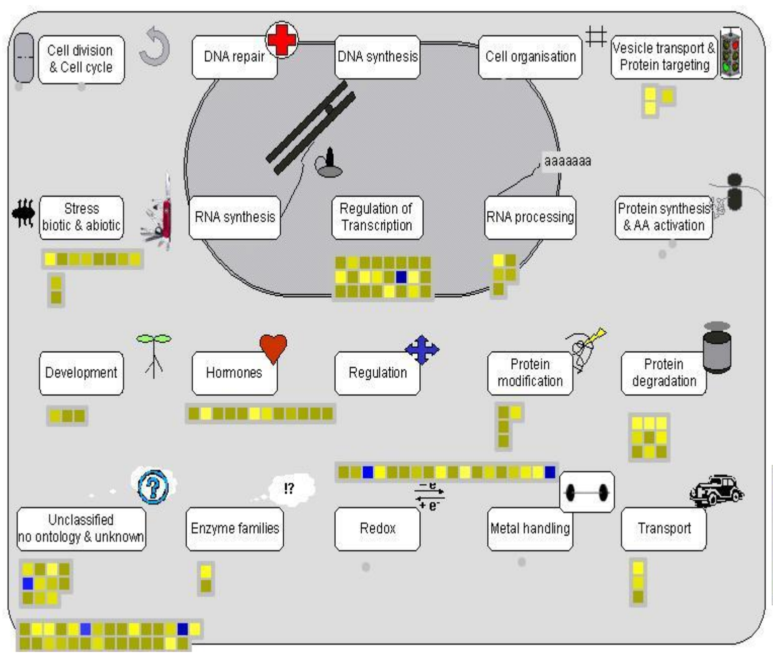

a. Transgene effect in controls | shoot | 3Hr | Cell function overview

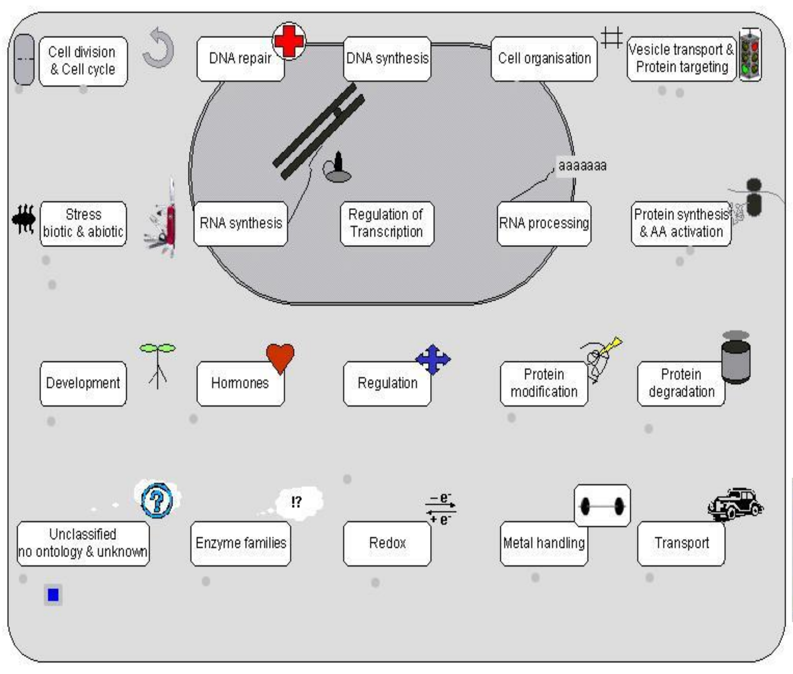

b. Transgene effect in controls| root | 51Hr | Cell function overview

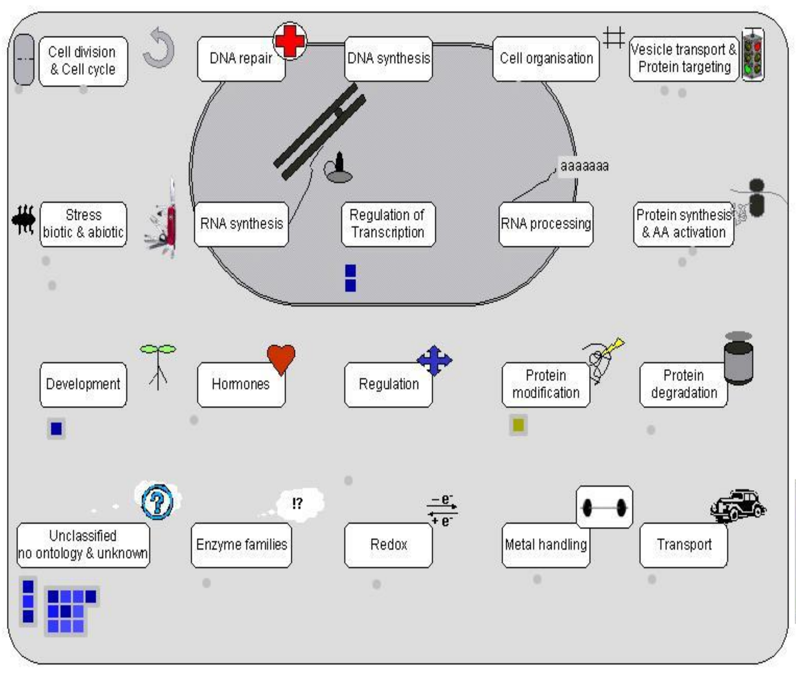

c. Transgene effect in controls | shoot | 51Hr | Cell function overview

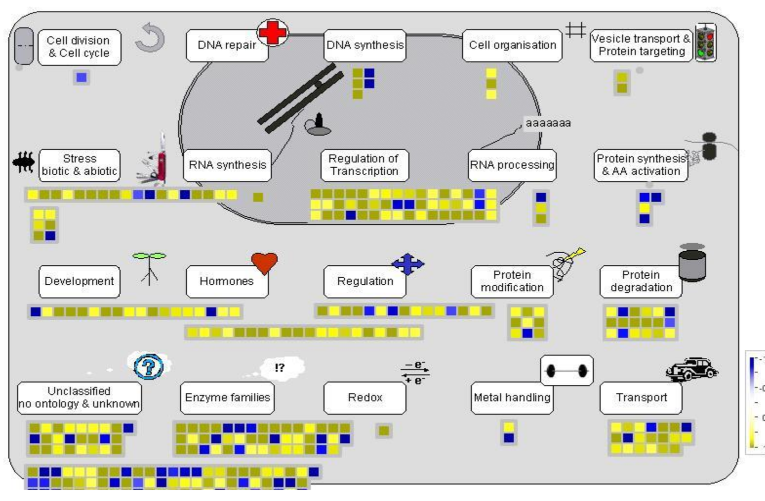

d. Transgene effect in salt | root | 3Hr | Cell function overview

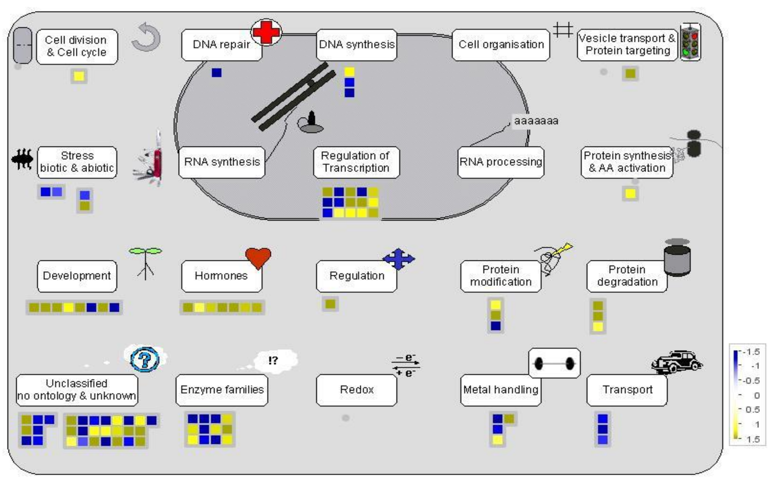

e. Transgene effect in salt | shoot | 3Hr | Cell function overview

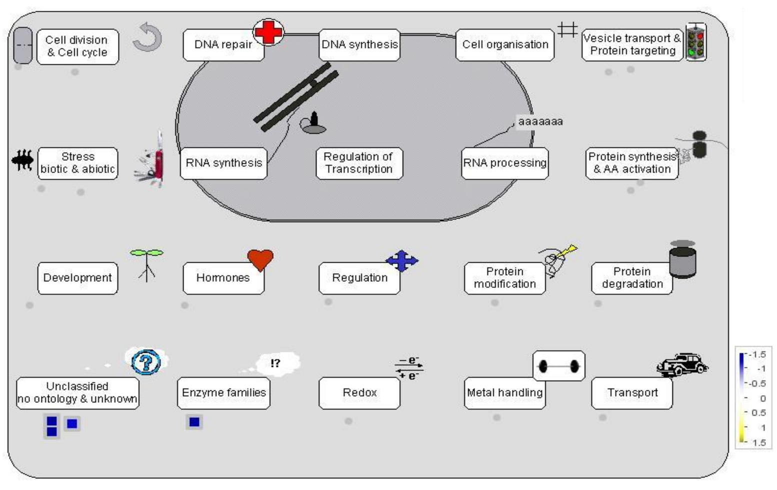

f. Transgene effect in salt | root | 51Hr | Cell function overview

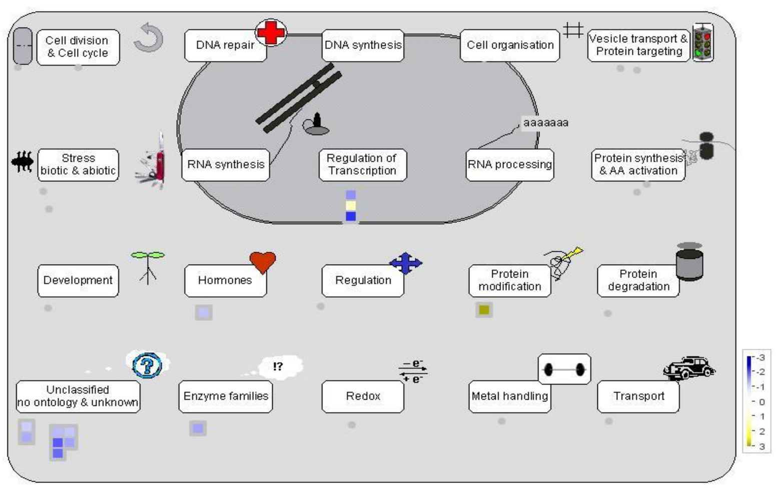

g. Transgene effect in salt | shoot | 51Hr | Cell function overview

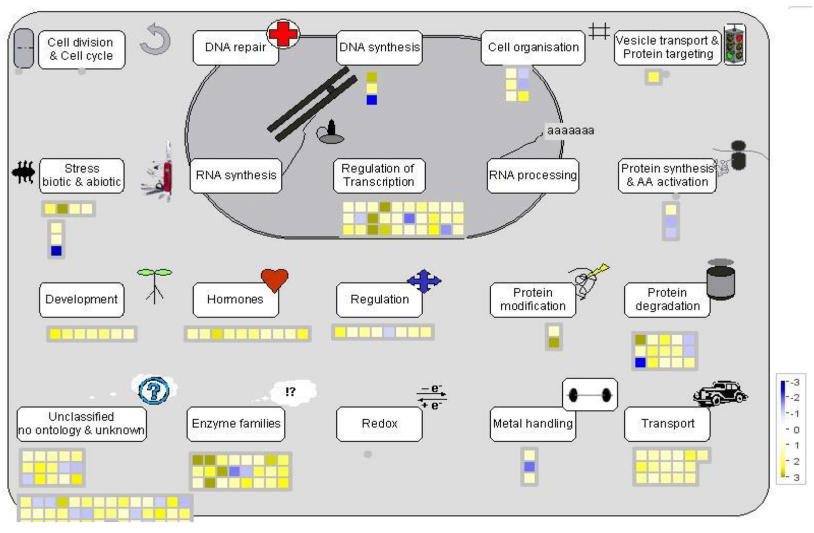

h. Transgene dependent salt response | root | 3Hr | Cell function overview

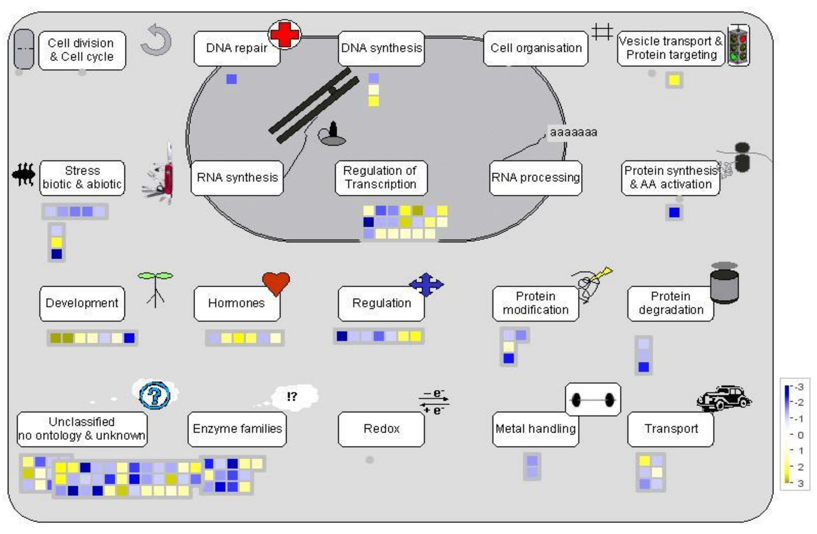

i. Transgene dependent salt response | shoot | 3Hr | Cell function overview

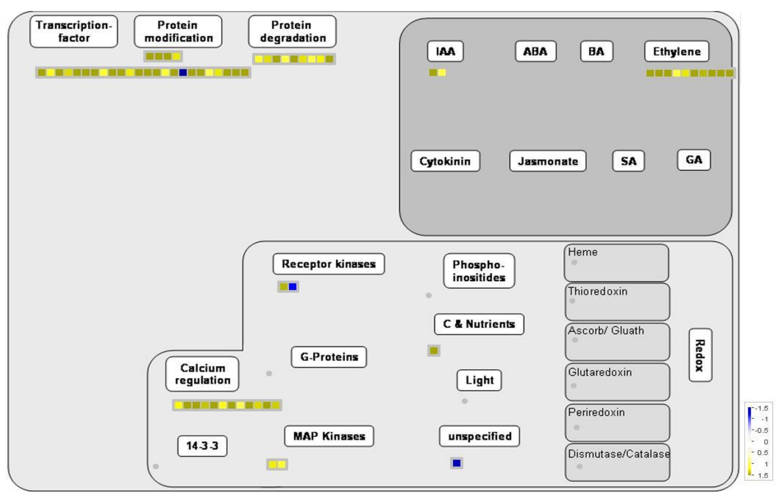

j. Transgene effect in controls | shoot | 3Hr | Regulation overview

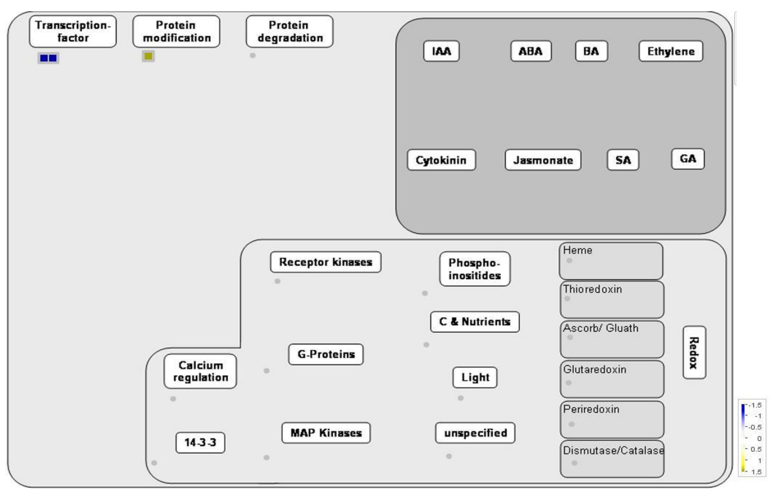

k. Transgene effect in controls | shoot | 51Hr | Regulation overview

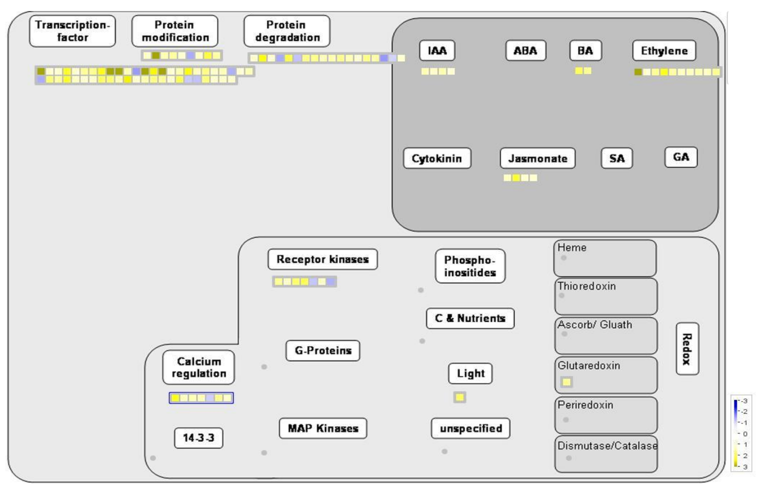

l. Transgene effect in salt | root | 3Hr | Regulation overview

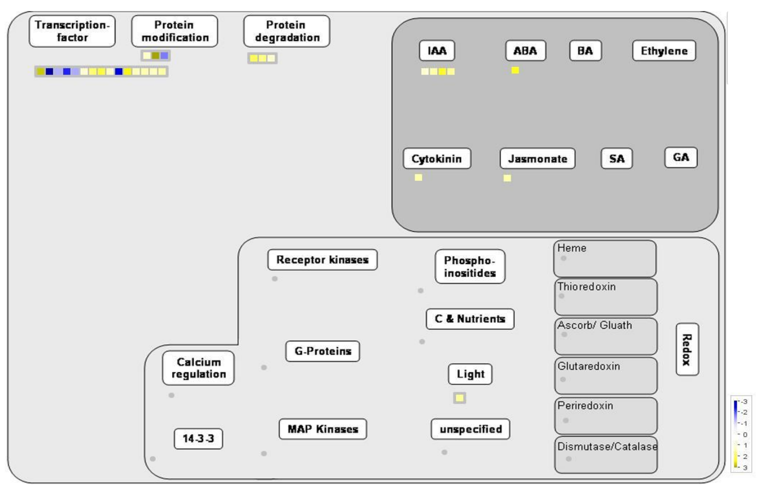

m Transgene effect in salt | shoot | 3Hr | Regulation overview

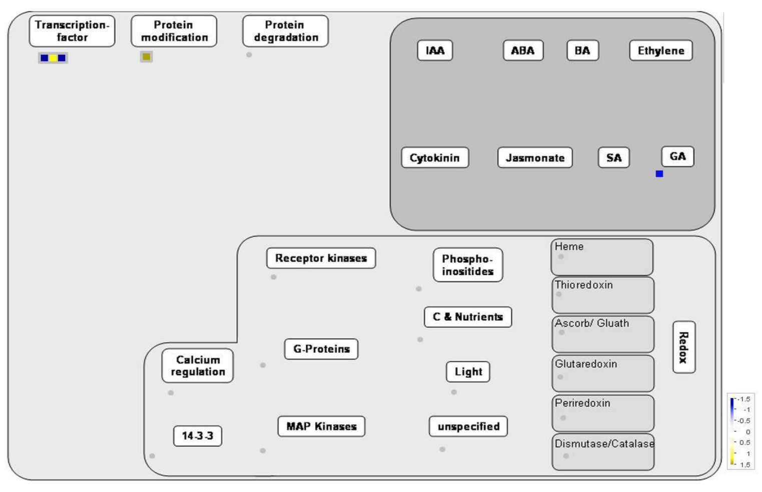

n. Transgene effect in salt | shoot | 51Hr | Regulation overview

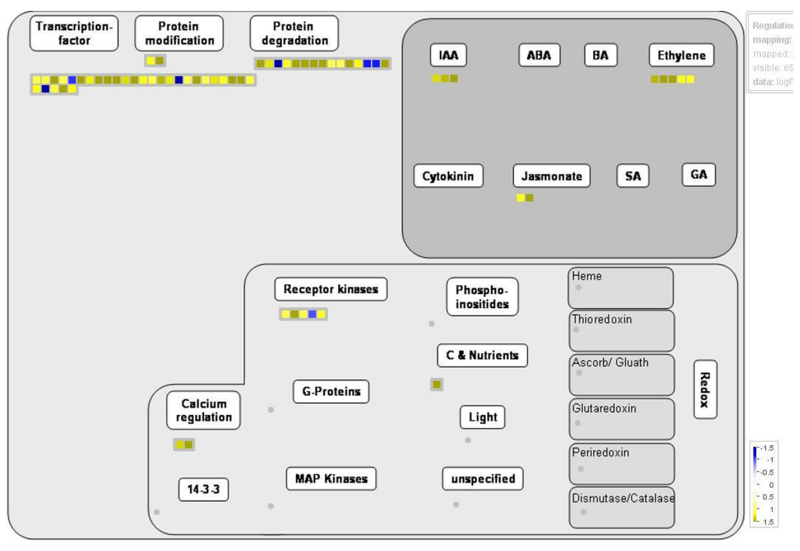

o. Transgene dependent salt response | root | 3Hr | Regulation overview

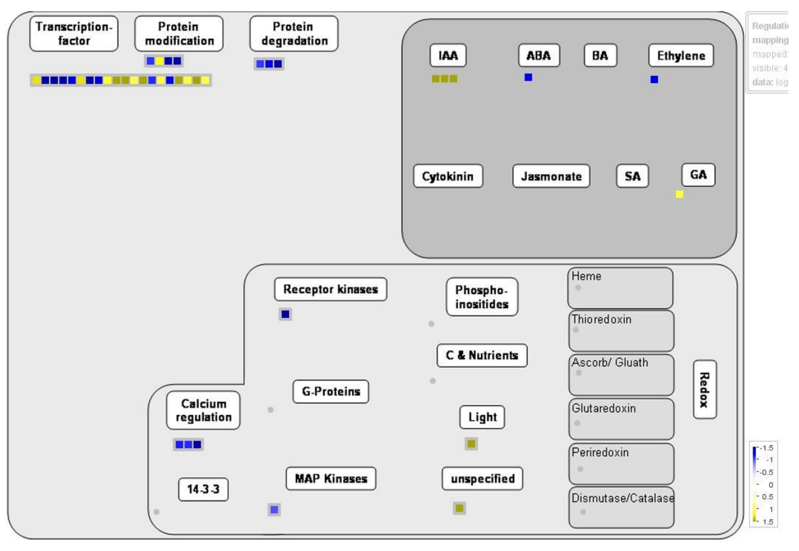

p. Transgene dependent salt response | shoot | 3Hr | Regulation overview

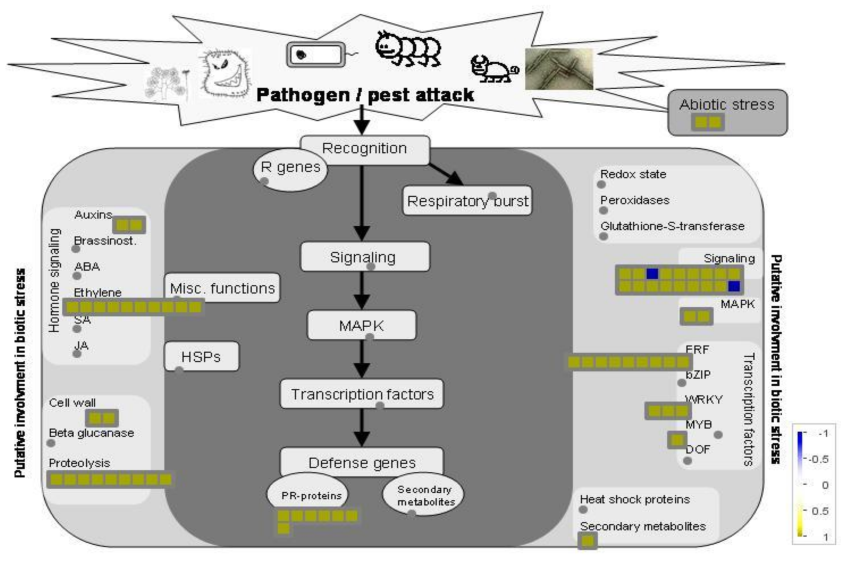

q. Transgene effect in controls | shoot | 3Hr | Putative Biotic stress pathways

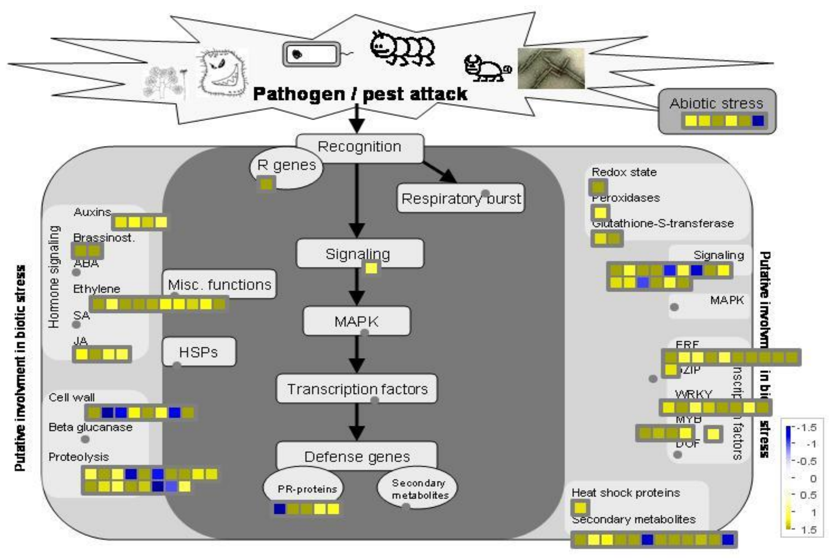

r. Transgene effect in salt | root | 3Hr | Putative Biotic stress pathways

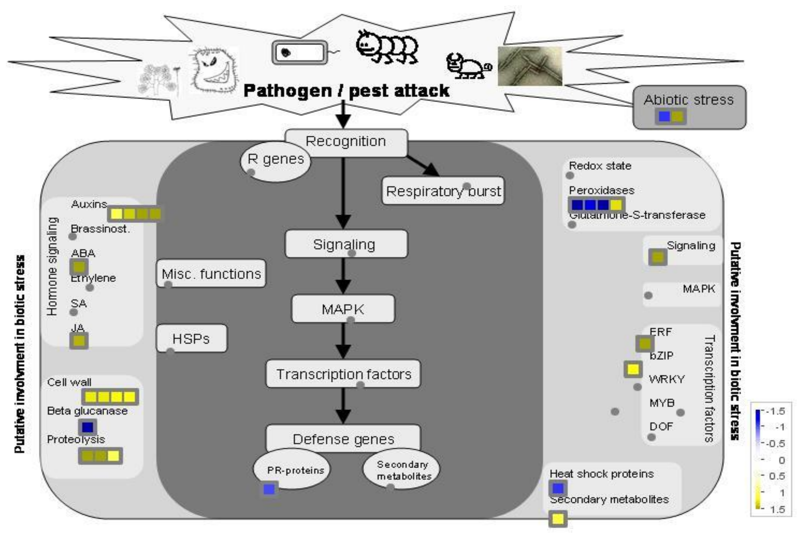

s. Transgene effect in salt | shoot | 3Hr | Putative Biotic stress pathways

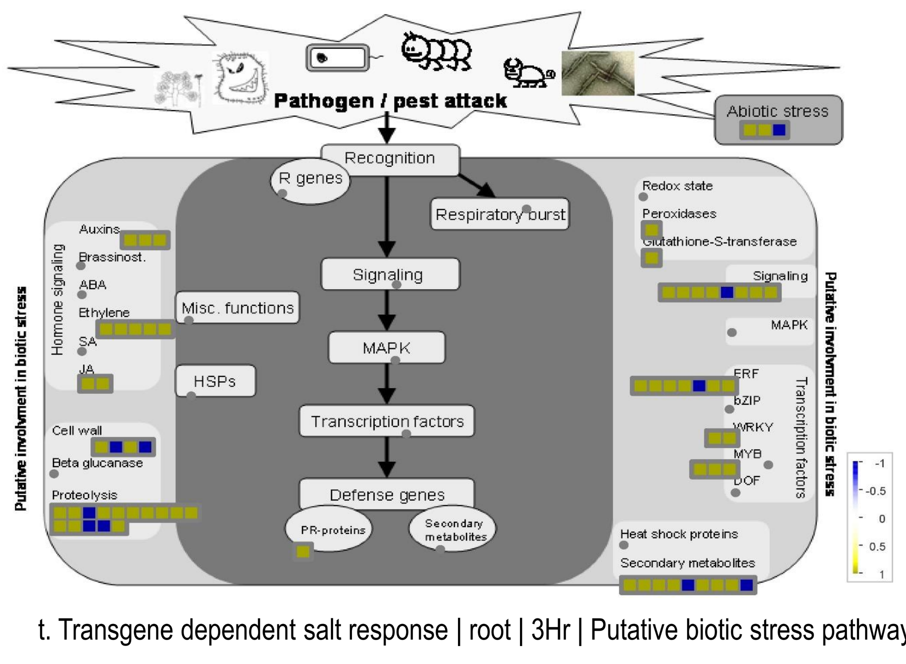

t. Transgene dependent salt response | root | 3Hr | Putative biotic stress pathways

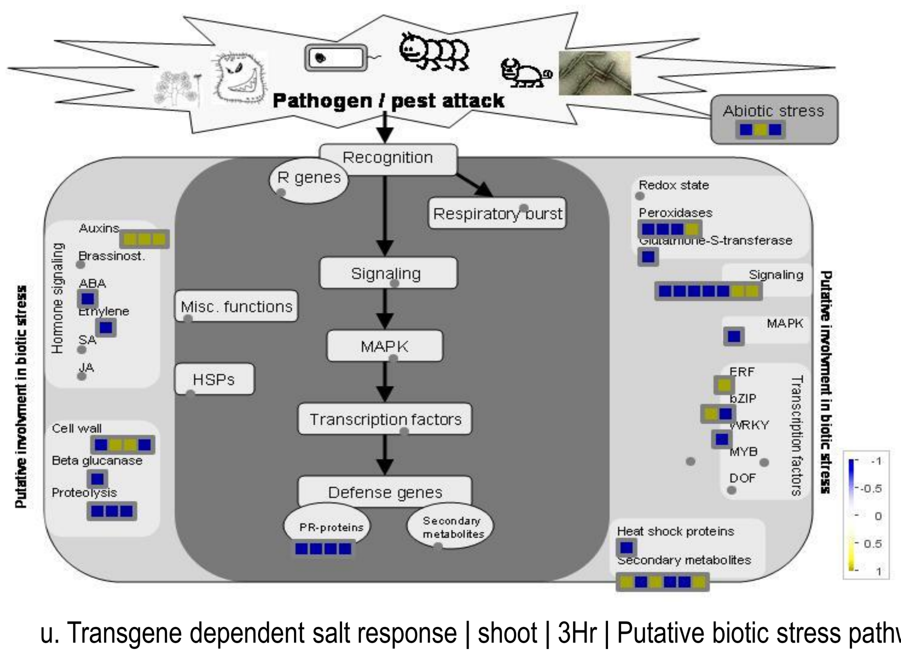

u. Transgene dependent salt response | shoot | 3Hr | Putative biotic stress pathways

### S8. MapMan pathway analysis of the DEGs

a-i : cell function overview ; j-p: regulation overview ; q-u: putative biotic stress pathways
