## Supplementary material for "AtCIPK16 Mediates Salt Stress Through Phytohormones and Transcription Factors": S9

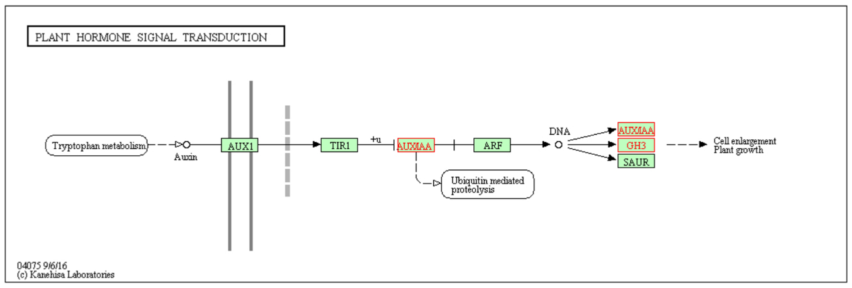


a) Transgene Effect in Salt | Root | 3Hr | Transport related genes in hormone signal transduction pathway

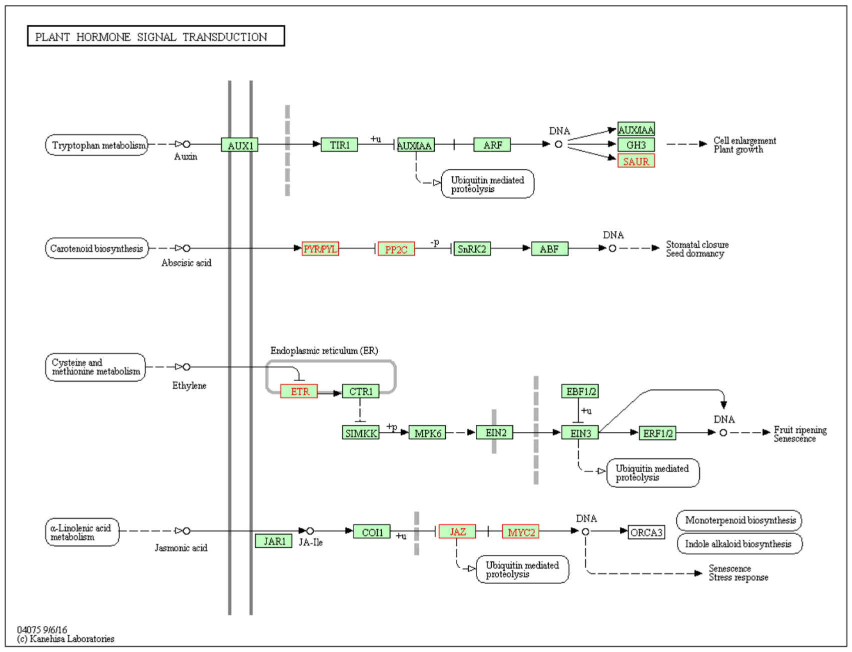


b) Transgene Dependent Salt Response | Root | 3Hr | Transcription factors involved in hormone signal transduction pathway

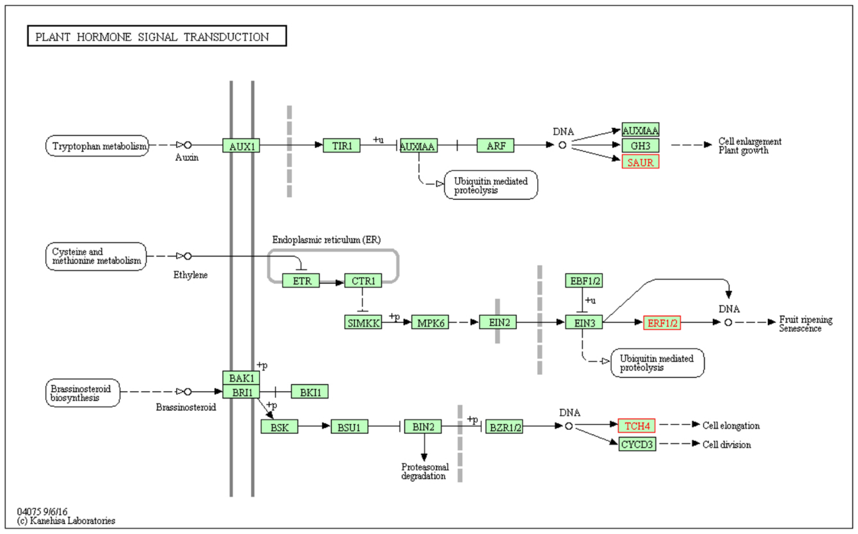


c) Transgene Effect in Controls| Shoot | 3Hr | Hormone metabolism genes in hormone signal transduction pathway

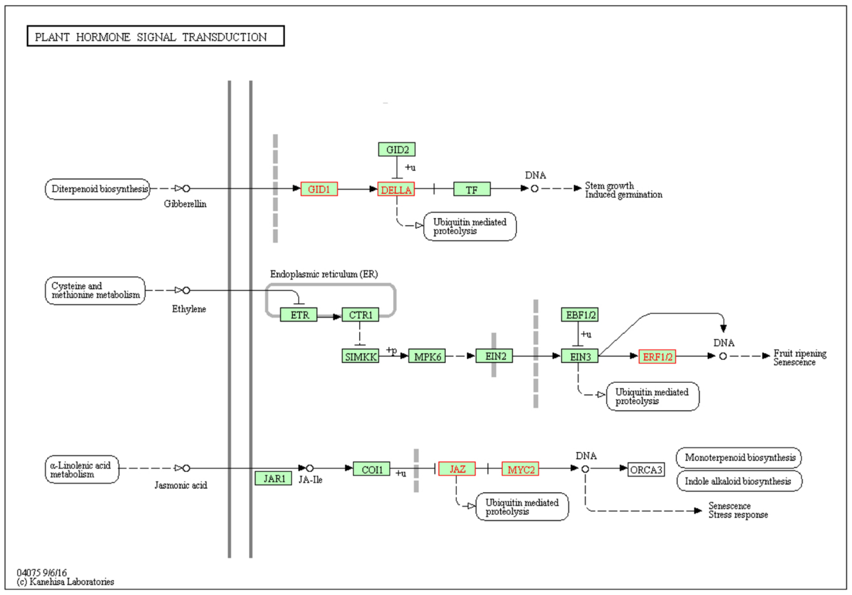


d) Transgene Effect in Salt | Root | 3Hr | Hormone metabolism genes in hormone signal transduction pathway

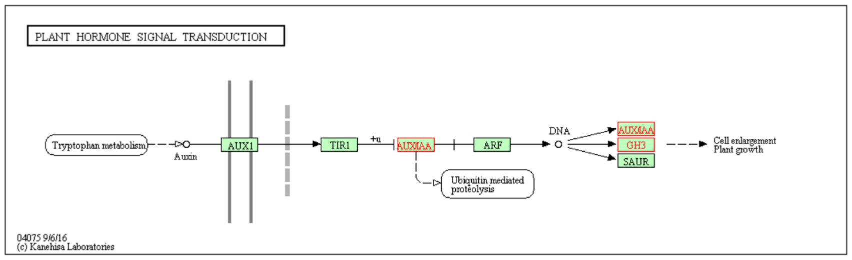


e) Transgene Effect in Salt | Shoot | 3Hr | Hormone metabolism genes in hormone signal transduction pathway

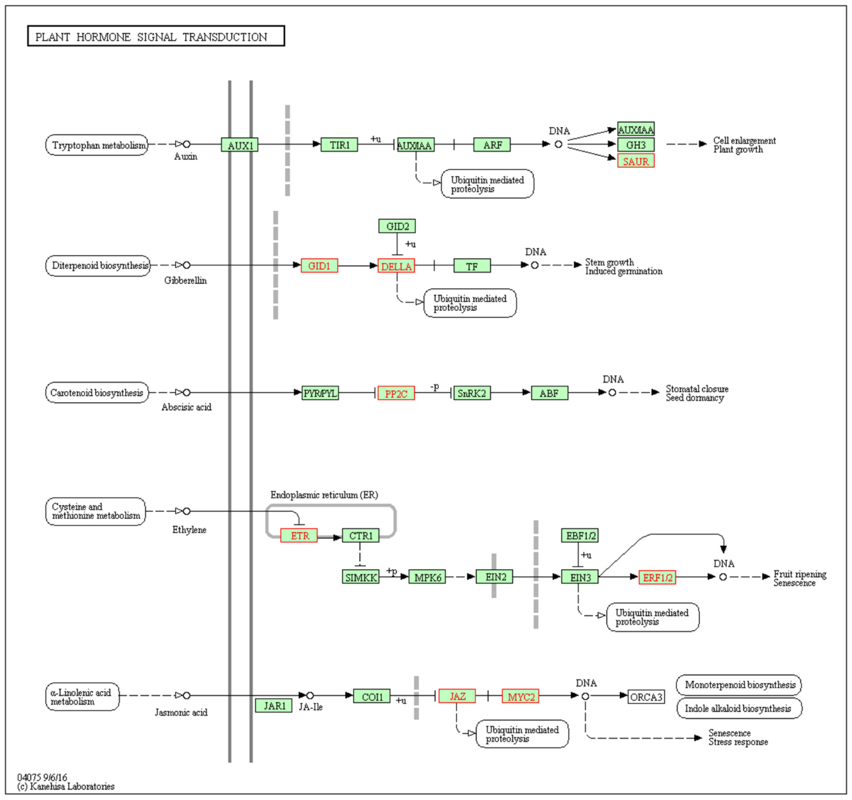


f) Transgene Dependent Salt Response | Root | 3Hr | Hormone metabolism genes in hormone signal transduction pathway

S9: KEGG Pathways of DEG subsets that are discussed in Mapman categories and their associated genes identified throughATTED-II.
The pathways are autogenerated through the search and colour pathway foption in Kyoto Encyclopaedia of Genes and Genomes (KEGG) server (www.genome.jp/kegg/) The input genes for each section are the up regulated DEGs for a), c), d) and e). and all DEGs for b) and f). The associated genes that are included in the list are retrieved throough the NetworkDrawer tool with default options (Platform is automatic, CoEx option add many genes and PPI option add a few genes) of ATTED-II server (atted.jp/). Rectangular boxes with green colour background represent genes in the pathway, arrows represent a molecular interaction or a relationship . The red framed green boxes with red letters show the genes that are in the input lists, each group of DEGs used to mine the pathways in seach sub figure are mentioned below the respective sub figure a)-f), empty circles represent chemical molecules, rrectangles with rounded edges lshows the ink from the current pathway to another pathway, doubled lines represent the plasma membrane , the dashed grey lines are shown when a direct association between ttwo moleules are unknown.
