## Supplementary material for "AtCIPK16 Mediates Salt Stress Through Phytohormones and Transcription Factors": S10

**S10. KEGG pathways enriched for the DEG subsets of selected MapMan categories and their associated genes identified through ATTED-II**

a) Transporters

Transgene Effect in Salt | Root | 3Hr

| KEGG* ID | Title | #genes |
|---|---|---|
| ath01110 | Biosynthesis of secondary metabolites | 16 |
| ath00940 | Phenylpropanoid biosynthesis | 9 |
| ath04626 | Plant-pathogen interaction | 7 |
| ath00360 | Phenylalanine metabolism | 7 |
| ath04075 | Plant hormone signal transduction | 4 |

Transgene Dependent Salt Response | Root | 3Hr

| KEGG* ID | Title | #genes |
|---|---|---|
| ath01110 | Biosynthesis of secondary metabolites | 15 |
| ath00360 | Phenylalanine metabolism | 5 |
| ath04075 | Plant hormone signal transduction | 4 |
| ath00280 | Valine, leucine and isoleucine degradation | 4 |
| ath04626 | Plant-pathogen interaction | 4 |

Transgene Dependent Salt Response | Shoot | 3Hr

| KEGG* ID | Title | #genes |
|---|---|---|
| ath01110 | Biosynthesis of secondary metabolites | 3 |
| ath00592 | alpha-Linolenic acid metabolism | 3 |
| ath00040 | Pentose and glucuronate interconversions | 2 |
| ath00073 | Cutin, suberine and wax biosynthesis | 2 |

b) Regulation of transcription

Transgene Effect in Controls | Shoot | 3Hr

| KEGG* ID | Title | #genes |
|---|---|---|
| ath04626 | Plant-pathogen interaction | 11 |
| ath01110 | Biosynthesis of secondary metabolites | 3 |
| ath04075 | Plant hormone signal transduction | 3 |
| ath03018 | RNA degradation | 3 |
| ath00270 | Cysteine and methionine metabolism | 2 |

Transgene Effect in Salt| Root | 3Hr

| KEGG* ID | Title | #genes |
|---|---|---|
| ath04626 | Plant-pathogen interaction | 3 |
| ath04075 | Plant hormone signal transduction | 2 |

Transgene Effect in Salt| Shoot | 3Hr

| KEGG* ID | Title | #genes |
|---|---|---|
| ath01110 | Biosynthesis of secondary metabolites | 4 |
| ath00903 | Limonene and pinene degradation | 2 |
| ath04120 | Ubiquitin mediated proteolysis | 2 |
| ath00500 | Starch and sucrose metabolism | 2 |
| ath00945 | Stilbenoid, diarylheptanoid and gingerol biosynthesis | 2 |

Transgene Dependent Salt Response | Root | 3Hr

| KEGG* ID | Title | #genes |
|---|---|---|
| ath04626 | Plant-pathogen interaction | 17 |
| ath01110 | Biosynthesis of secondary metabolites | 15 |
| ath04075 | Plant hormone signal transduction | 14 |
| ath00500 | Starch and sucrose metabolism | 7 |
| ath00940 | Phenylpropanoid biosynthesis | 5 |

Transgene Dependent Salt Response | Shoot | 3Hr

| KEGG* ID | Title | #genes |
|---|---|---|
| ath01110 | Biosynthesis of secondary metabolites | 3 |
| ath04075 | Plant hormone signal transduction | 3 |
| ath00280 | Valine, leucine and isoleucine degradation | 3 |
| ath04626 | Plant-pathogen interaction | 2 |
| ath00500 | Starch and sucrose metabolism | 2 |

c) Metal synthesis and assimilation related

Transgene Effect in Salt | Root | 3Hr

| EGG* ID | Title | #genes |
|---|---|---|
| ath01110 | Biosynthesis of secondary metabolites | 2 |
| ath00480 | Glutathione metabolism | 2 |

Transgene Effect in Salt | Shoot | 3Hr

| KEGG* ID | Title | #genes |
|---|---|---|
| ath00860 | Porphyrin and chlorophyll metabolism | 3 |
| ath01110 | Biosynthesis of secondary metabolites | 2 |

Transgene Dependent Salt Response | Root | 3Hr

| KEGG* ID | Title | #genes |
|---|---|---|
| ath01110 | Biosynthesis of secondary metabolites | 3 |
| ath00860 | Porphyrin and chlorophyll metabolism | 3 |
| ath01230 | Biosynthesis of amino acids | 2 |
| ath00480 | Glutathione metabolism | 2 |
| ath01200 | Carbon metabolism | 2 |

d) Enzyme families

Transgene Effect in Salt | Root | 3Hr

| KEGG* ID | Title | #genes |
|---|---|---|
| ath01110 | Biosynthesis of secondary metabolites | 28 |
| ath00480 | Glutathione metabolism | 10 |
| ath00945 | Stilbenoid, diarylheptanoid and gingerol biosynthesis | 10 |
| ath00903 | Limonene and pinene degradation | 10 |
| ath01130 | Biosynthesis of antibiotics | 7 |

Transgene Effect in Salt | Shoot | 3Hr

| KEGG* ID | Title | #genes |
|---|---|---|
| ath01110 | Biosynthesis of secondary metabolites | 7 |
| ath00360 | Phenylalanine metabolism | 6 |
| ath00940 | Phenylpropanoid biosynthesis | 6 |
| ath00073 | Cutin, suberine and wax biosynthesis | 3 |
| ath04146 | Peroxisome | 2 |

Transgene Dependent Salt Response | Root | 3Hr

| KEGG* ID | Title | #genes |
|---|---|---|
| ath01110 | Biosynthesis of secondary metabolites | 9 |
| ath00480 | Glutathione metabolism | 4 |
| ath00940 | Phenylpropanoid biosynthesis | 4 |
| ath00061 | Fatty acid biosynthesis | 3 |
| ath04075 | Plant hormone signal transduction | 3 |

Transgene Dependent Salt Response | Shoot | 3Hr

| KEGG* ID | Title | #genes |
|---|---|---|
| ath01110 | Biosynthesis of secondary metabolites | 17 |
| ath00360 | Phenylalanine metabolism | 6 |
| ath00940 | Phenylpropanoid biosynthesis | 6 |
| ath00941 | Flavonoid biosynthesis | 5 |
| ath00520 | Amino sugar and nucleotide sugar metabolism | 2 |

f) Hormone metabolism

Transgene Effect in Controls | Shoot | 3Hr

| KEGG* ID | Title | #genes |
|---|---|---|
| ath04626 | Plant-pathogen interaction | 9 |
| ath04075 | Plant hormone signal transduction | 9 |
| ath01110 | Biosynthesis of secondary metabolites | 3 |
| ath03018 | RNA degradation | 3 |
| ath00270 | Cysteine and methionine metabolism | 2 |

Transgene Effect in Salt | Root | 3Hr

| KEGG* ID | Title | #genes |
|---|---|---|
| ath01110 | Biosynthesis of secondary metabolites | 16 |
| ath04075 | Plant hormone signal transduction | 10 |
| ath04626 | Plant-pathogen interaction | 9 |
| ath00592 | alpha-Linolenic acid metabolism | 7 |
| ath01130 | Biosynthesis of antibiotics | 3 |

Transgene Effect in Salt | Shoot | 3Hr

| KEGG* ID | Title | #genes |
|---|---|---|
| ath04075 | Plant hormone signal transduction | 6 |
| ath01110 | Biosynthesis of secondary metabolites | 3 |
| ath00500 | Starch and sucrose metabolism | 3 |
| ath00908 | Zeatin biosynthesis | 2 |
| ath00942 | Anthocyanin biosynthesis | 2 |

Transgene Dependent Salt Response | Root | 3Hr

| KEGG* ID | Title | #genes |
|---|---|---|
| ath01110 | Biosynthesis of secondary metabolites | 17 |
| ath04075 | Plant hormone signal transduction | 16 |
| ath04626 | Plant-pathogen interaction | 13 |
| ath00592 | alpha-Linolenic acid metabolism | 10 |
| ath00591 | Linoleic acid metabolism | 4 |

Transgene Dependent Salt Response | Shoot | 3Hr

| KEGG* ID | Title | #genes |
|---|---|---|
| ath01110 | Biosynthesis of secondary metabolites | 5 |
| ath00360 | Phenylalanine metabolism | 3 |
| ath00280 | Valine, leucine and isoleucine degradation | 3 |
| ath00940 | Phenylpropanoid biosynthesis | 3 |
| ath00942 | Anthocyanin biosynthesis | 2 |

g) Putative biotic stress related signalling pathways

Transgene Effect in Controls | Shoot | 3Hr

| KEGG* ID | Title | #genes |
|---|---|---|
| ath04626 | Plant-pathogen interaction | 14 |
| ath01110 | Biosynthesis of secondary metabolites | 6 |
| ath04130 | SNARE interactions in vesicular transport | 3 |
| ath04075 | Plant hormone signal transduction | 3 |
| ath00564 | Glycerophospholipid metabolism | 3 |

Transgene Effect in Salt | Root | 3Hr

| KEGG* ID | Title | #genes |
|---|---|---|
| ath04626 | Plant-pathogen interaction | 6 |
| ath04075 | Plant hormone signal transduction | 5 |

Transgene Effect in Salt | Shoot | 3Hr

| KEGG* ID | Title | #genes |
|---|---|---|
| ath00500 | Starch and sucrose metabolism | 2 |

Transgene Dependent Salt Response | Root | 3Hr

| KEGG* ID | Title | #genes |
|---|---|---|
| ath04626 | Plant-pathogen interaction | 5 |
| ath01110 | Biosynthesis of secondary metabolites | 4 |
| ath00360 | Phenylalanine metabolism | 3 |
| ath00940 | Phenylpropanoid biosynthesis | 3 |

Transgene Dependent Salt Response | Shoot | 3Hr

| KEGG* ID | Title | #genes |
|---|---|---|
| ath04626 | Plant-pathogen interaction | 4 |
| ath01110 | Biosynthesis of secondary metabolites | 3 |
| ath00500 | Starch and sucrose metabolism | 2 |
| ath00940 | Phenylpropanoid biosynthesis | 2 |
